# LEA motifs promote desiccation tolerance *in vivo*

**DOI:** 10.1101/2021.02.04.429810

**Authors:** Jonathan D. Hibshman, Bob Goldstein

## Abstract

**Background:** Cells and organisms typically cannot survive in the absence of water. However, there are some notable exceptions, including animals such as nematodes, tardigrades, rotifers, and some arthropods. One class of proteins known to play a role in desiccation resistance is the late embryogenesis abundant (LEA) proteins. These largely disordered proteins protect plants and animals from desiccation. A multitude of studies have characterized stress-protective capabilities of LEA proteins *in vitro* and in heterologous systems. However, the extent to which LEA proteins exhibit such functions *in vivo*, in their native contexts in animals, is unclear.

Furthermore, little is known about the distribution of LEA proteins in multicellular organisms or tissue-specific requirements in conferring stress protection.

**Results:** To study the endogenous function of an LEA protein in an animal, we created a true null mutant of *C. elegans* LEA-1, as well as endogenous fluorescent reporters of the protein. We confirmed that *C. elegans* lacking LEA-1 are sensitive to desiccation. LEA-1 mutant animals were also sensitive to heat and osmotic stress and were prone to protein aggregation. During desiccation, LEA-1 expression increased and became more widespread throughout the body. LEA-1 was required at high levels in body wall muscle for animals to survive desiccation and osmotic stress. We identified minimal motifs within *C. elegans* LEA-1 that are sufficient to increase desiccation survival of *E. coli*. To test whether such motifs are central to LEA-1’s *in vivo* functions, we then replaced the sequence of *lea-1* with these minimal motifs and found that *C. elegans* survived mild desiccation and osmotic stress at the same levels as worms with the full-length protein.

**Conclusions:** Our results provide insights into the endogenous functions and expression dynamics of an LEA protein in a multicellular animal. The results show that LEA-1 buffers animals from a broad range of stresses. Our identification of LEA motifs that can function in both bacteria and in a multicellular organism suggests the possibility of engineering LEA-1-derived peptides for optimized desiccation protection.

## Background

Animals regularly encounter abiotic stresses due to fluctuating environments. One such stress is desiccation. Water is essential for cellular metabolism, and without it cellular components including DNA, RNA, proteins, and membranes are unstable. However, certain animals can lose nearly all their internal water and yet survive (anhydrobiosis). This group includes nematodes, rotifers, tardigrades, and certain arthropods such as some crustaceans and insects. Studying the mechanisms by which these organisms are able to survive desiccation is of fundamental interest and may lead to better understanding of biological protectants.

One family of common protectants with demonstrated efficacy during desiccation is the late embryogenesis abundant (LEA) proteins. These proteins were originally identified in cotton seeds and were subsequently found to be present in many other plants [1–6]. LEA proteins protect the viability of desiccated seeds [7]. Animal LEA proteins were later identified in the nematode *Aphelenchus avenae* and other anhydrobiotic animals [8–13]. Several families of LEA proteins have been described, each containing distinct motifs [14, 15]. Nematode LEA proteins both in *A. avenae* as well as the more common model system *C. elegans* are members of the Group 3 LEA proteins. This is the most common LEA variety within anhydrobiotic animals [16]. LEA proteins are thought to be intrinsically disordered, i.e. largely unstructured in hydrated conditions, but under water-limiting conditions the proteins acquire recognizable alpha-helical secondary structure [17–19]. There are several examples in which heterologous expression of LEA proteins confers stress resistance in a number of host cell types, including yeast, bacteria, rice, and mammalian cell culture [20–25]. Additionally, isolated LEA protein has been demonstrated to protect against protein aggregation and stabilize membranes and liposomes *in vitro* [26–28]. Although the functions of LEA proteins are well-established, most studies have utilized heterologous expression systems. Therefore, we have limited knowledge of the endogenous, *in vivo* roles of LEA proteins.

The nematode *C. elegans* is an ideal model for the study of the endogenous functions of LEA proteins. *C. elegans* survive many stresses including desiccation, have a well-resourced genetic toolkit, and contain two LEA family proteins, *lea-1* and *dur-1* (<u>d</u>auer <u>u</u>p-regulated) [10, 29]. Disruption of *lea-1* has been reported to sensitize worms to desiccation, osmotic stress with sucrose, and larval heat stress [10, 29, 30]. *dur-1* is also required for a robust desiccation response but is apparently unable to compensate for the loss of LEA-1 [29]. Whereas most studies show that heterologous expression is sufficient to protect against stresses in other systems, these *C. elegans* studies provide evidence for necessity in the native function of LEA proteins in animals. However, little is known about the tissues in which LEA proteins are required or the mechanisms by which they function *in vivo* [14, 31, 32].

We used LEA-1 of *C. elegans* as a model to study the endogenous function of an LEA protein in a multicellular animal. We confirmed and expanded phenotypes associated with disruption of LEA-1, characterized expression patterns and dynamics in response to desiccation, and identified minimal amino acid motifs within the protein that harbor the biochemical properties to confer desiccation resistance. These results provide insights into the *in vivo*, endogenous functions of an LEA protein in a multicellular animal.

## Results

### Generation of a null allele of *lea-1*

We were motivated to determine the role of LEA-1 in the endogenous organismal context of *C. elegans*. Previous studies have relied on RNAi for gene knockdown and did not fully eliminate *lea-1* mRNA [10] or noted that RNAi seldom fully eliminates the target mRNA [29]. One recent study used a deletion allele, but only confirmed the phenotype of desiccation sensitivity [30]. Furthermore, there are many isoforms of LEA-1 (Fig. 1), and the potential for alternative isoform usage makes mutant analysis difficult to interpret. To address these concerns, we created a full-length, 15.8 kb deletion of the *lea-1* locus via CRISPR, and we also cleaned the genetic background of a second, pre-existing mutant allele, *lea-1(tm6452)*, by outcrossing it to wild-type worms for 5 generations [10, 29, 33, 34]. For the full-length deletion (*lea-1Δ*), the deleted gene was replaced with a *pmyo-2*::GFP::*myo-2 3’-UTR* construct as a reporter for the deletion (Fig. 1). The successful deletion was confirmed via visualization of pharyngeal GFP as well as by PCR (Fig. S1A-D).

**Figure 1.**
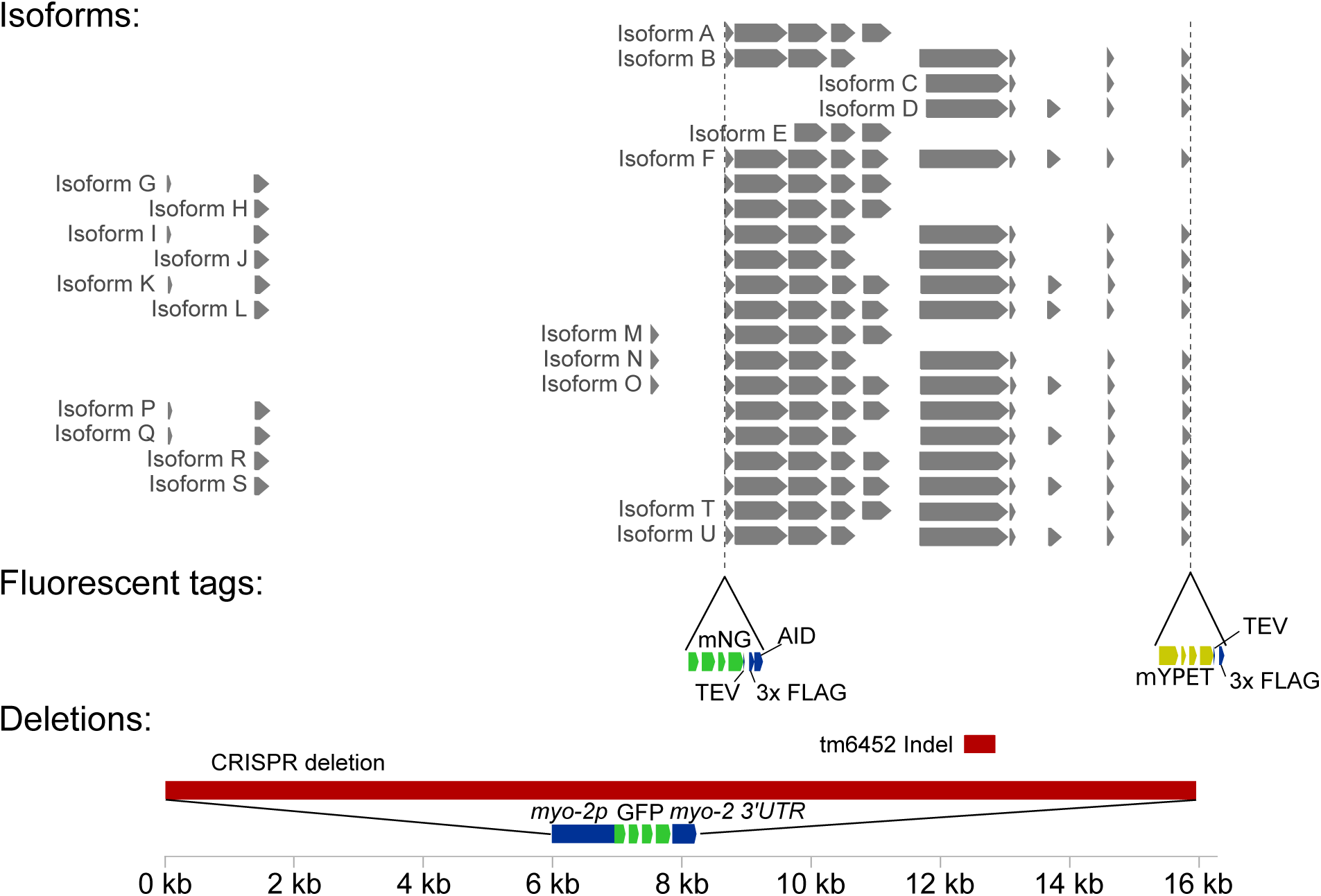
Predicted isoform structure of LEA-1 and generation of a null allele and two endogenous fluorescent tags. Isoform annotations were combined from the UCSC genome browser (http://genome.ucsc.edu) from the Feb. 2013 release (WBcel235/ce11) and Wormbase version WS278. Locations for the existing tm6452 indel mutation, the newly created CRISPR null mutation, and the insertion of mNeonGreen and mYPET fluorescent tags are indicated.

### *lea-1* deletion mutants are sensitive to multiple stresses

LEA proteins are most well known for promoting survival during desiccation. Because worms are reported to survive desiccation only when in the dauer state [35], and *daf-2(e1370)* offers a temperature-sensitive constitutive dauer phenotype when worms are grown at 25 °C [36], we chose to conduct the majority of our experiments in this *daf-2* background to facilitate consistent production of dauer worms. In a *daf-2(e1370)* background, we confirmed that both *lea-1Δ* and *lea-1(tm6452)* dauer worms were sensitive to desiccation at high relative humidity (97.5%) and at a more stringent desiccation stress of 60% humidity (Fig. 2A), consistent with published results [10, 29, 30]. Additionally, dauer worms of each of these mutant strains were sensitive to osmotic stress in a concentrated salt solution. While there were no differences in survival in worms kept in water for 2 hr, both *lea-1Δ* and *lea-1(tm6452)* worms had significantly lower survival when exposed to 1M NaCl for 2 hr (Fig. 2B). Both mutants were also sensitive to exposure to heat shock at 37 °C (Fig. 2C).

**Figure 2.**
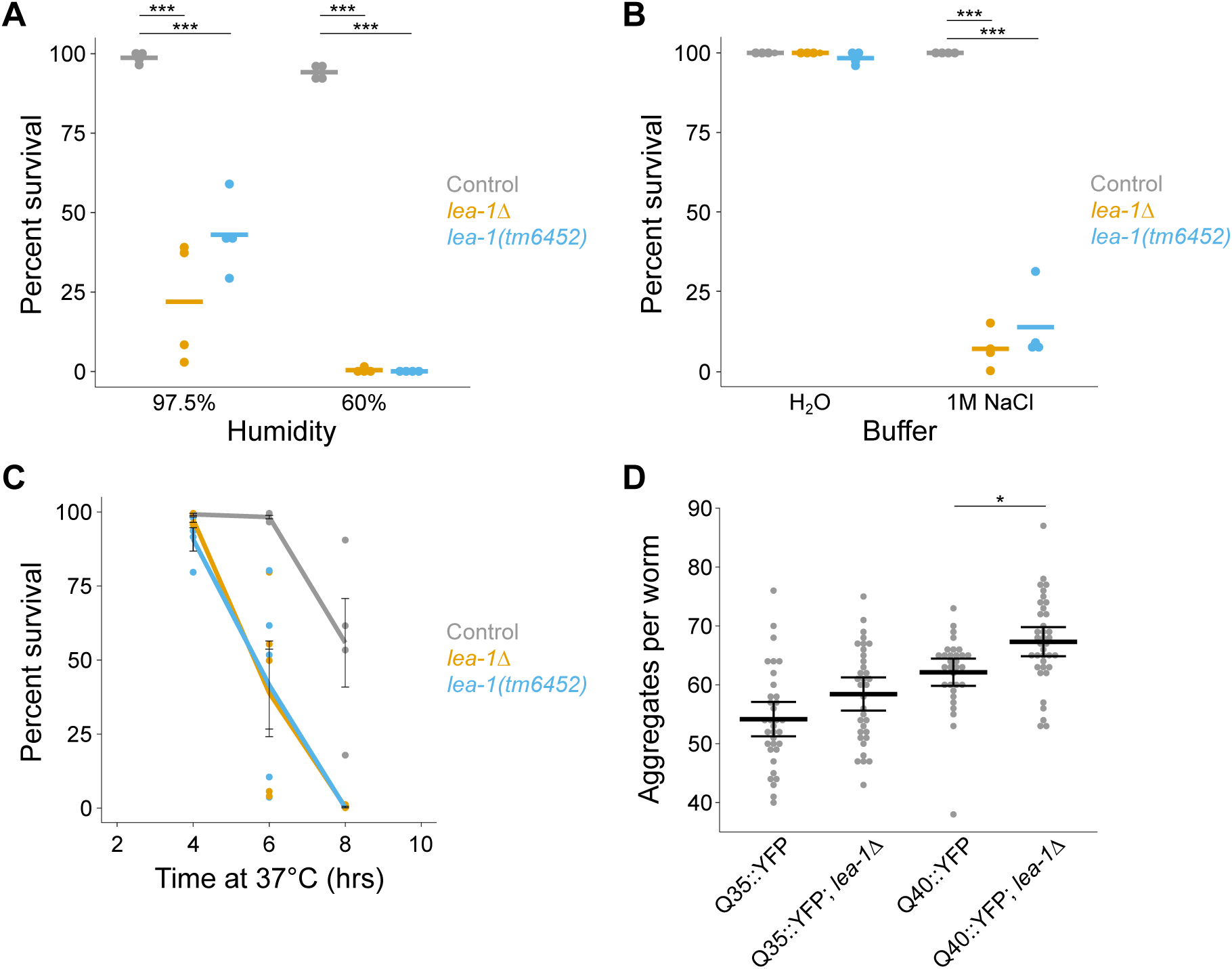
*lea-1* is required for tolerance of desiccation and osmotic stress. **A)** Dauer larvae of *lea-1Δ* and *lea-1(tm6452)* mutants are sensitive to desiccation. Survival is significantly reduced in worms exposed to 97.5% relative humidity for 4 days (p=0.0002, p=0.0001 respectively, n=4, unpaired T-test) as well as 60% for 1 day after 4 days at 97.5% (p=2.5×10^-10^, p=1.8×10^-10^ respectively, n=4, unpaired T-test). Data points represent results from individual experiments and bars represent mean survival. **B)** Dauer larvae of *lea-1Δ* and *lea-1(tm6452)* mutants are sensitive to osmotic stress in 1M NaCl for 2 hr (p=8.8×10^-8^, p=6.1×10^-6^ respectively, n=4, unpaired T-test). Data points represent results from individual experiments and bars represent mean survival. **C)** Dauer larvae of *lea-1Δ* and *lea-1(tm6452)* mutants are sensitive to heat stress at 37°C (p<0.0001 for each mutant allele relative to control, n=4-5 replicates per timepoint, 2- way ANOVA). Lines depict mean survival and error bars represent SEM. Data points depict survival from individual experiments. **D)** *lea-1Δ* mutants have a higher number of polyglutamine protein aggregates. The total number of aggregates in the body wall muscle of individual worms from three independent biological replicates is shown. Thick bars indicate the mean number of aggregates and error bars depict 95% confidence intervals. Worms carrying a transgene expressing a 35-glutamine repeat (Q35) with a YFP for visualization have only marginally more aggregates than controls (p=0.22, n=3, unpaired T-test), while worms carrying a Q40::YFP transgene have significantly more aggregates than controls (p=0.02, n=3, unpaired T-test). Note, all worms were in a *daf-2(e1370)* background to facilitate dauer formation at 25°C. * indicates p<0.05, *** indicates p<0.001.

LEA proteins have been demonstrated to prevent protein aggregation *in vitro* and in cell culture [26, 37–39]. To test for *in vivo* roles in preventing protein aggregation, we assessed the number of polyglutamine aggregates in body wall muscle of *lea-1Δ* animals. These animals expressed tracts of 35 or 40 glutamines fused to a yellow fluorescent protein (YFP) reporter [40]. Mutants with the Q35::YFP reporter did not have a statistically significant difference from controls (p=0.22); however, *lea-1Δ* animals with a Q40::YFP reporter had significantly more aggregates per worm than controls (p=0.02) after 5 days of development and dauer formation (Fig. 2D). We conclude that LEA-1 has *in vivo* functions protecting animals against desiccation, osmotic stress, heat stress, and formation of polyglutamine protein aggregates.

### LEA-1 lacks discernable nonredundant functions outside of stress tolerance

Working with a true null mutant allowed us to ask if there were any phenotypes associated with deletion of *lea-1* in the context of normal development and physiology. We tested effects on lifespan and brood size as especially sensitive quantitative proxies for diverse effects on physiology or development. Our results suggest that LEA-1 functions specifically in response to stress: In a wild-type (N2) background we did not observe any significant differences in lifespan or brood size of *lea-1Δ* animals (Fig. 3A,B). We also did not observe significant differences in polyglutamine aggregation during aging in these worms when maintained under normal laboratory conditions (Fig. S2). These results indicate that LEA-1 may function primarily during the dauer stage and in response to conditions of external stress.

**Figure 3.**
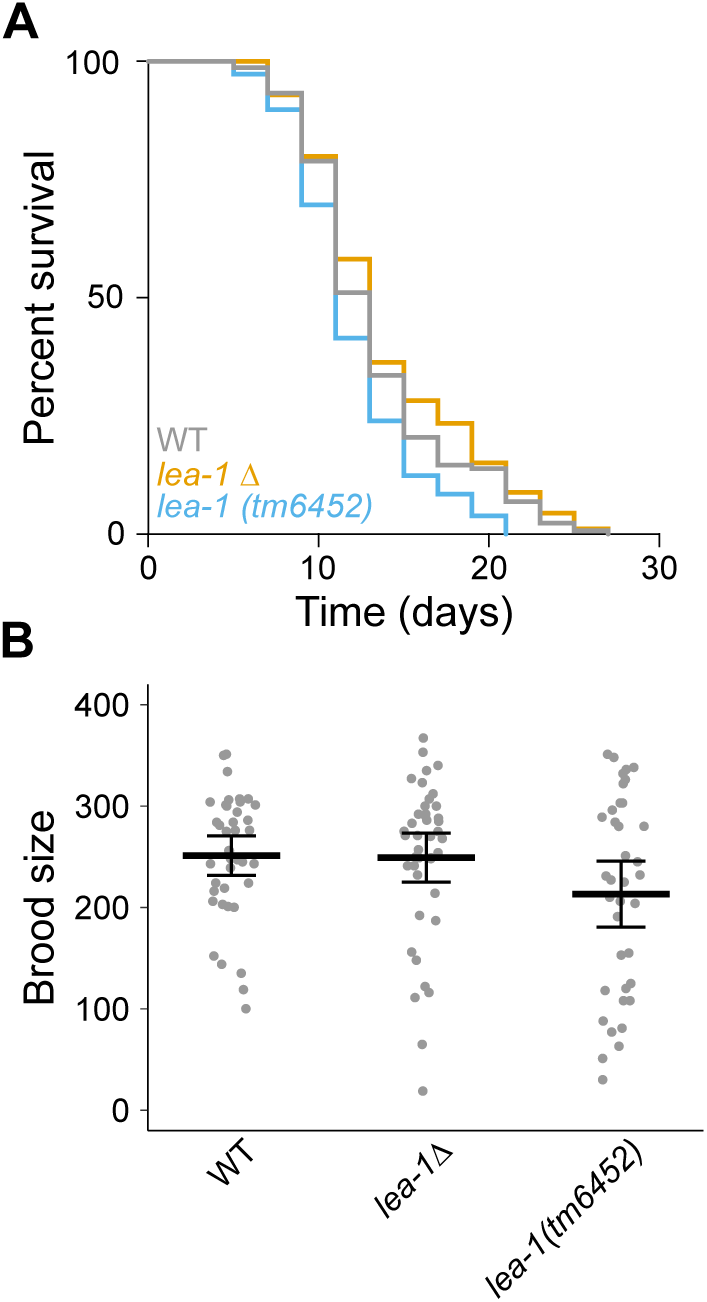
Phenotypes of LEA-1 are specific to conditions of stress. **A)** Lifespan is not significantly altered in either *lea-1Δ* or *lea-1(tm6452)* mutants in a WT (N2) background. **B)** There are no significant differences in brood size of *lea-1Δ* (p=0.97, n=3, unpaired T-test) or *lea- 1(tm6452)* mutants (p=0.41, n=3, unpaired T-test) relative to N2 worms. Brood size of individual worms from independent replicates are shown. The mean is also shown and error bars represent 95% confidence intervals. Lifespan and brood size experiments were carried out in a WT (N2) background.

### Expression of multiple LEA-1 isoforms is increased in response to desiccation

To examine where LEA-1 is expressed *in vivo*, we used fluorescent monomeric NeonGreen (mNG) and monomeric yellow fluorescent protein (mYPET) tags to label endogenous LEA-1 at two different positions [41, 42]. Sequence for mNG as well as a 3x FLAG and auxin inducible degron sequence were inserted in the N-terminal region of the gene (Fig. 1). Similarly, sequence for mYPET and 3x FLAG were added to the C-terminus of LEA-1 (Fig. 1). As with the *lea-1Δ* mutant, these edits were confirmed via visualization of fluorescence and PCR (Fig. S1E). When desiccated, animals expressing the fluorescent tags survived equally as well as *daf-2* control animals (Fig. 4A). We also confirmed that the fluorescent tags did not significantly impact osmotic stress survival in 1M NaCl for 2 hr (Fig. S3A). We conclude that introduction of either fluorophore did not disrupt functions of the endogenous LEA-1 protein.

**Figure 4.**
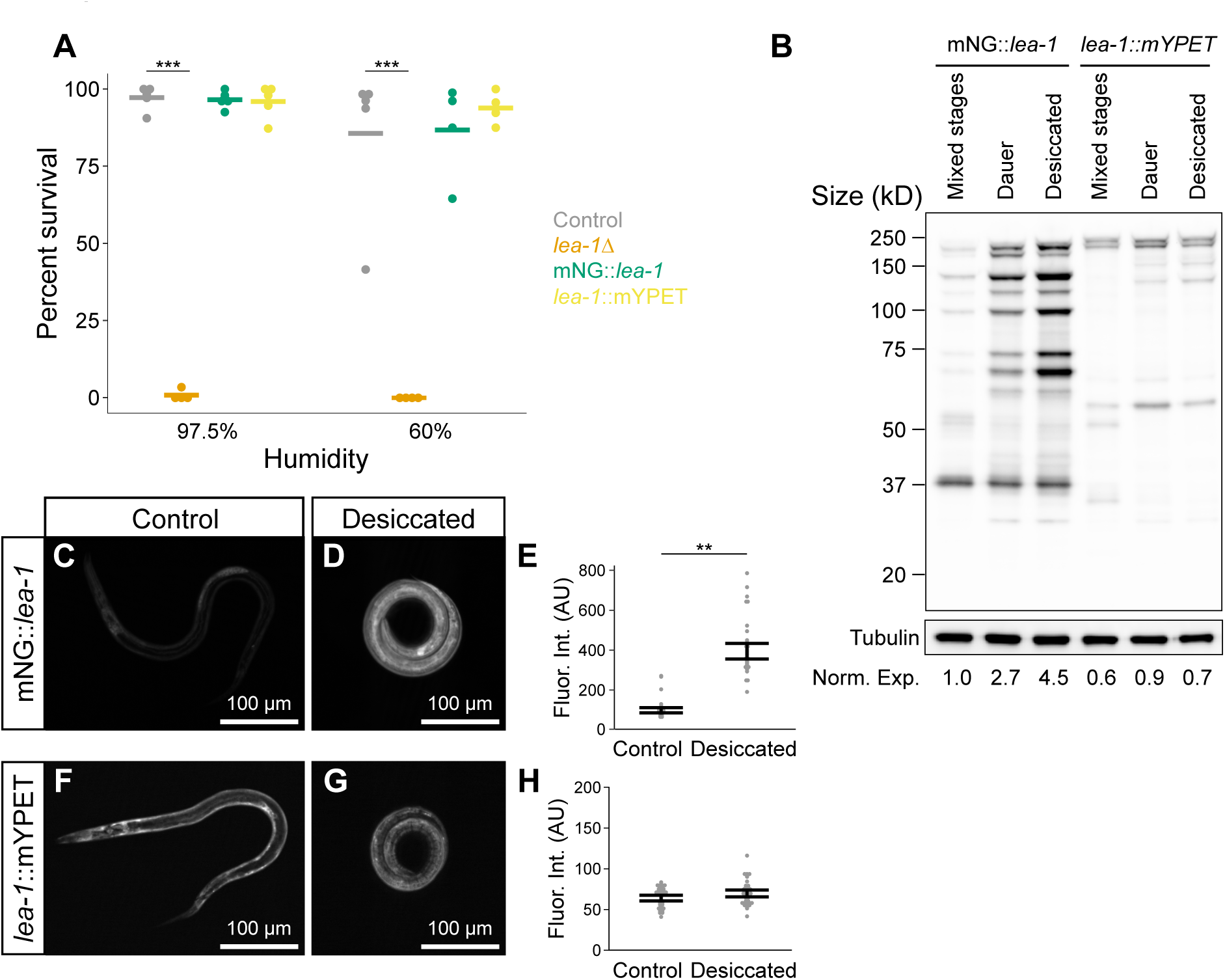
Functional endogenous fluorescent tags reveal increased LEA-1 protein expression of multiple isoforms during desiccation. **A)** Endogenous fluorescent tags do not disrupt the function of LEA-1 in desiccation survival. *lea-1Δ* are sensitive to desiccation at 97.5% relative humidity (p=6.6×10^-10^, n=4, unpaired T-test) and 60% relative humidity (p=0.0002, n=4, unpaired T-test). Survival of mNeonGreen tagged LEA-1 worms (mNG::*lea-1*) and mYPET tagged worms (*lea- 1*::mYPET) was not significantly different from *daf-2(e1370)* controls after exposure to 97.5% or 60% RH. **B)** Protein samples extracted from of mixed stage cultures, dauer worms, and desiccated worms were blotted with an anti-FLAG antibody indicates multiple isoforms tagged by the two independent tags. The membrane was stripped and blotted for tubulin as a loading control. Total protein in each lane was quantified and normalized to the tubulin loading control. Normalized proteins expression measurements, relative to mixed stage mNG::*lea-1* animals, are listed at the bottom of each lane (Norm. Exp.). **C)** A representative dauer worm expressing mNG::*lea-1*. **D)** A representative desiccated dauer larvae expressing mNG::*lea-1*. **E)** mNG fluorescence is significantly increased in desiccated worms relative to non-desiccated controls (p=0.001, n=3 replicates with 14-15 worms per condition, unpaired T-test). **F)** A representative dauer worm expressing *lea-1*::mYPET. **G)** A desiccated dauer expressing *lea-1*::mYPET. **H)** Fluorescent intensity is not significantly different between control and desiccated dauer worms expressing *lea-1*::mYPET (p=0.39, n=3 replicates with 15 worms per condition, unpaired T-test). Note that all worms were in a *daf-2(e1370)* background. ** indicates p<0.01, *** indicates p<0.001.

LEA-1 protein is known to be upregulated in response to desiccation [10, 29]. However, there are many predicted isoforms of LEA-1 (Fig 1), and it is unclear whether all isoforms are upregulated or whether all of LEA-1’s protective roles involve such upregulation. To examine the diversity of isoforms that are actively expressed in different conditions we lysed cultures of mixed stage worms, dauer worms, and desiccated dauers to blot with an anti-FLAG antibody to recognize the 3x FLAG epitope appended to each fluorophore. Numerous isoforms appeared tagged with the 5’ mNG tag, while fewer isoforms of distinct sizes carrying the 3’ mYPET tag were prominent (Fig. 3B). Isoforms labeled with the 5’ mNG tag but not the 3’ mYPET tag were found to be upregulated during desiccation. These changes in expression were confirmed by quantification of *in vivo* fluorescence intensity in dauer and desiccated worms. Fluorescence from the mNG reporter was significantly increased after 4 days of desiccation at 97.5% relative humidity (Fig. 4C-E). Expression of mNG::LEA-1 was diffuse throughout the body during desiccation (Fig. 4D). In contrast, mYPET fluorescence level was not significantly altered during desiccation (Fig. 4F-H), consistent with the Western blot results. For both reporters, fluorescence intensity was not changed in response to 2 hr of osmotic shock in 1M NaCl (Fig. S3B-G). We conclude that only some isoforms of LEA-1 are upregulated in response to desiccation, and that *lea-1*’s response to osmotic shock does not appear to involve a similar upregulation.

### Depletion of LEA-1 in body wall muscle reduces survival of desiccation and osmotic stress

To examine the localization of upregulated LEA-1 isoforms in the stage when LEA-1 has a protective role, we visualized mNG::LEA-1 fluorescence in dauer larvae. We found that the protein was prominently expressed in specific tissues: the germline, body wall muscle, pharynx, and excretory cell (Fig. 5A-D). mNG::LEA-1 was localized to both the excretory cell body, which is adjacent to the pharynx (Fig. 5D), as well as excretory canals, which extend laterally along the length of the body (Fig. 5A). Some faint expression was seen in seam cells (Fig. 5B), as well as the intestine (Fig. 5C). Additionally, some unidentified cells in the head were fluorescent. The expression and localization of LEA-1 to particular tissues in dauer worms suggested that its function might be required in only some tissues or required at higher levels in some tissues than in others.

**Figure 5.**
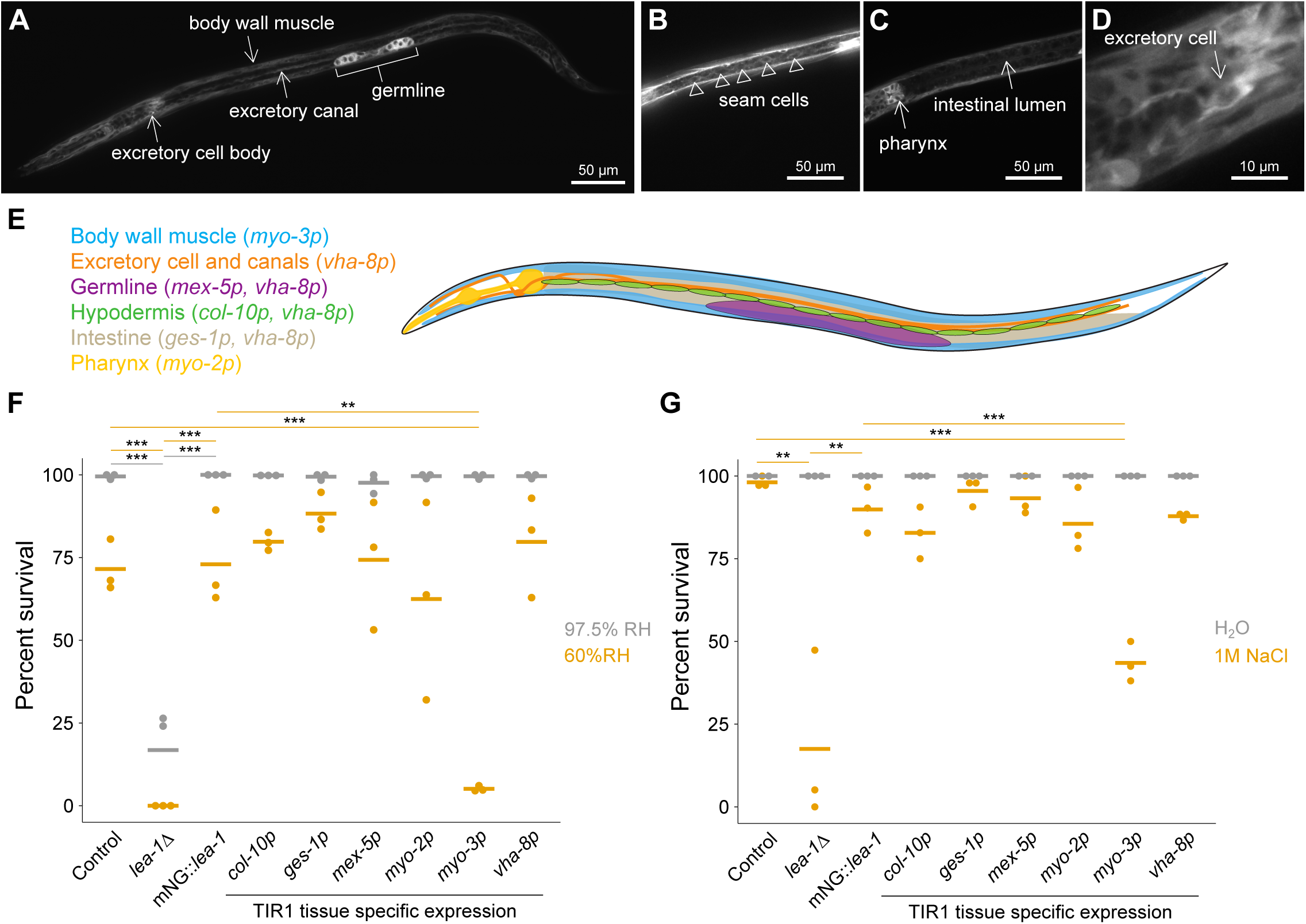
LEA-1 is required in body wall muscle to survive desiccation and osmotic stress. **A-D)** Representative images of an mNG::*lea-1* worm depict the major sites of expression. There is prominent fluorescence in the germline, body wall muscle, excretory cell, and pharynx. There is also some apparent expression in seam cells (B) and faint fluorescence in the intestine (C). A zoomed in image of the excretory cell is shown in (D). **E)** A cartoon depicts tissues in which LEA-1 is expressed that were targeted for protein depletion by driving TIR1 under the control of different promoters. **F)** Desiccation survival is plotted for worms with tissue-specific LEA-1 depletion by expression of TIR1 under various promoters and exposure to 1mM auxin. Depletion of the mNG::*lea-1* protein utilizing auxin-induced degradation reduces desiccation survival in worms expressing TIR1 under a *myo-3* (body wall muscle-specific) promoter relative to controls (p=0.0001, n=3, unpaired T-test) and worms expressing only mNG::*lea-1* and not TIR1 (p=0.001, n=3, unpaired T-test). No other site of TIR1 expression significantly altered desiccation survival at 60% RH. **G)** Survival of osmotic stress in 1M NaCl for 4 hrs is plotted for the same strains as in (E). *lea-1Δ* mutants and worms expressing TIR1 under a *myo-3* promoter were the only two strains that with significant differences in survival relative to both control and mNG::*lea-1*-expressing animals (*lea-1Δ*: vs. control p=0.006, vs. mNG::*lea-1* p=0.01; *myo-3p*: vs. control p=0.0001, vs. mNG::*lea-1* p=0.0009, unpaired T-tests, n=3). Note, the images in A-D show a worm with an N2 background, whereas worms in (E) and (F) are in a *daf-2(e1370)* background (control). ** indicates p<0.01, *** indicates p<0.001.

To determine the sites of action of LEA-1, we utilized auxin-induced degradation to deplete protein. We used the auxin-induced degron (AID) sequence that we included on the mNG::3xFLAG tag as a target for protein degradation when the modified F-box protein TIR1 is expressed and worms are exposed to auxin [43]. Expression of TIR1 under the control of different promoters allows for tissue-specific conditional depletion [44]. We used a panel of worms expressing TIR1 in the various tissues in which LEA-1 was expressed to determine the tissue or tissues in which LEA-1 was required for survival of desiccation and osmotic stress (Fig. 5E) [44]. These experiments were conducted in a *daf-2* mutant background, for which expression patterns of LEA-1 were not noticeably different from a WT (N2) background (Fig. S4A). When exposed to auxin, dauer worms showed depletion of mNG::LEA-1 in the expected locations for each promoter used: the intestine (using the *ges-1* promoter, called *ges-1*p), germline (*mex-5*p), pharynx (*myo-2*p), body wall muscle (*myo-3*p), and a combination of intestine, excretory cell and canal, germline, and some other cells in the head (*vha-8*p) (Fig. S4). Expression of TIR1 with a *col-10* promoter to target hypodermis did not seem to have a significant effect on expression, consistent with the apparently minimal baseline hypodermal expression of mNG::LEA-1 (Fig. S4C). Additionally, the *myo-2*p::TIR1 strain depleted some pharyngeal mNG::LEA-1 protein, but did not entirely eliminate expression, particularly in the posterior bulb of the pharynx (Fig. S4F).

When exposed to desiccation at 97.5% relative humidity, no TIR1 expression strain had significantly reduced survival relative to controls (*daf-2*) or worms expressing mNG::LEA-1 alone (Fig. 5F). However, when exposed to 60% relative humidity, *myo-3*p::TIR1 worms had significantly reduced survival. This suggests that LEA-1 is required in the body wall muscle to promote desiccation resistance. When exposed to osmotic stress with 1M NaCl for 4 hrs, we observed a similar result. *lea-1Δ* mutants and *myo-3*p::TIR1 worms had significantly reduced survival compared to control (*daf-2*) and mNG::LEA-1-expressing worms (Fig. 5G). Collectively, these results suggest that LEA-1 synthesized in the body wall muscle is required for survival of multiple stresses, and that other tissues do not require high levels of the tagged LEA-1 isoforms for survival of animals.

### LEA-1 minimal motifs confer desiccation resistance to bacteria

Having described *in vivo* roles for an animal LEA protein and its localized expression and function, we sought to identify the protein’s protective domains. Given the variety of isoforms of *lea-1* in *C. elegans* that are expressed during desiccation (Fig. 1, Fig. 4B), before testing specific domains *in vivo* we employed a heterologous expression approach to efficiently test if different isoforms improved desiccation tolerance in *E. coli* equally, or if some were more effective than others. We selected isoforms A, D, E, F, and K for expression because they have a combination of overlapping and non-overlapping sequence to allow for deduction of regions of interest in the case that some isoforms are more effective at conferring desiccation resistance (Fig. 6A). We confirmed robust expression of each of the 5 different bacterially codon-optimized isoforms in BL21 *E. coli* and found that expression of each isoform conferred stress resistance to the bacteria, relative to cells expressing GFP (Fig. 6B, Fig. S5). Expression of a truncated version of GFP or actin (*C. elegans act-2*) did not increase desiccation resistance of bacteria (Fig. 6B).

**Figure 6.**
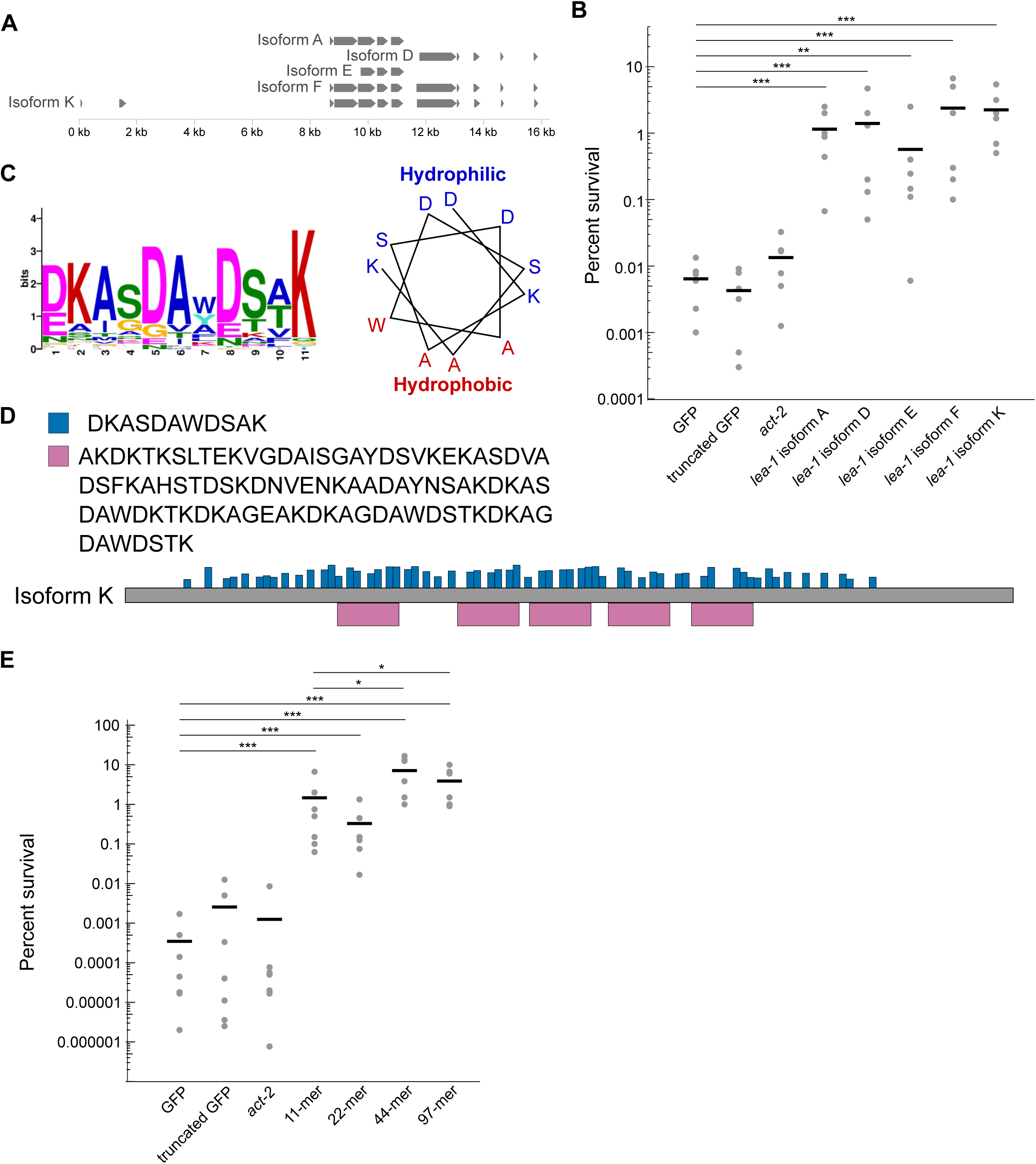
Heterologous expression of *C. elegans lea-1* isoforms and motifs improves bacterial desiccation survival. **A)** Select isoforms of LEA-1 used for bacterial expression. **B)** Desiccation survival of BL21 *E. coli* expressing codon-optimized *C. elegans* LEA-1 isoforms. Heterologous expression of each isoform increased desiccation survival (Isoform A p=2×10^-5^, Isoform D p=0.0002, Isoform E p=0.003, Isoform F p=9×10^-5^, Isoform K p=7×10^-7^, unpaired T-tests vs. GFP, n=6). **C)** A consensus 11-mer motif found in LEA-1 likely forms an amphipathic alpha helix. The position weight matrix of amino acids in the motif is shown, as well as a wheel diagram depicting the relative position of each consensus amino acid in an alpha helical conformation. **D)** Consensus amino acid sequences are shown for the 11-mer as well as a 97- mer motif that was also detected with an expanded motif window size. The frequency and distribution of occurrences of these motifs within isoform K are shown. **E)** Desiccation survival of BL21 *E. coli* is significantly increased by expression of motifs of LEA-1. Codon-optimized sequences for the 11-mer, as well as concatenated repeats of the 11-mer sequence (2x=22- mer, 4x=44-mer), and the 97-mer motif were expressed. Expression of each peptide increased survival of desiccated bacteria (11-mer p=2×10^-6^, 22-mer p=5×10^-6^, 44-mer p=1×10^-7^, 97-mer p=1×10^-7^, unpaired T-tests vs. GFP, n=7).

Because all five isoforms tested were sufficient to improve bacterial desiccation resistance, we hypothesized that motifs shared by these isoforms might be sufficient for conferring bacterial desiccation resistance. Group 3 LEA proteins commonly have repeated motifs of 11 amino acids that are capable of protecting heterologous cells against stresses including low pH, high salt, and desiccation-induced protein aggregation [8, 21, 39, 45–48]. These motifs are predicted to form alpha helices, and circular dichroism of LEA proteins and motif-containing peptides has confirmed alpha helix formation during conditions when water is limiting [17, 18, 49]. We used MEME Suite to detect a repeated 11-mer from the protein sequence of one of the longest *C. elegans* isoforms, LEA-1K [50]. An 11-mer was found that is predicted to form an amphipathic alpha-helix (Fig. 6C). When search parameters were relaxed to include motifs of other lengths, a 97-mer peptide encompassing multiple 11-mers was also identified. In the 1397 amino acid protein, there were five occurrences of the 97-mer and 61 instances of the 11-mer (Fig. 6D). To determine if these motifs alone from *C. elegans* LEA-1 were sufficient to improve desiccation tolerance, we expressed codon-optimized versions of these idealized motifs in *E. coli*. We included the base 11-mer, repeats containing two (22-mer) and four (44-mer) copies of its sequence, and the 97-mer. Expression of each of these minimal motifs significantly improved desiccation survival of the bacteria (Fig. 6E). Thus, it is likely that any of LEA-1’s isoforms that contain such repeats would improve survival of desiccation.

### LEA-1 motifs are sufficient for desiccation tolerance and osmotic stress survival *in vivo*

Because motifs of LEA-1 improved desiccation survival of bacteria, we were motivated to test if minimal motifs of LEA-1 are sufficient for stress tolerance *in vivo*. To specifically determine if minimal LEA-1 motifs are sufficient for desiccation tolerance in *C. elegans*, we deleted nearly all possible exons of *lea-1* and replaced them with sequence encoding single codon-optimized 44-mer or 97-mer motifs fused to mNG to allow visual confirmation of expression (Fig. 7A). Genomic sequence encoding the most N-terminal predicted exons for some isoforms (like the short N-terminal exons of isoforms K and N as shown in Fig. 7A) was left intact because these regions likely overlap with the promoter region or other regulatory sequence. These short predicted exons do not encode any 11-mer motifs. We confirmed genomic edits by PCR genotyping (Fig. S6A,B). We also determined that the expression patterns of each inserted motif::mNG fusion are similar to the overall pattern observed for tagged versions of the endogenous protein, suggesting that these genomic edits did not significantly disrupt gene regulation (Fig. S6C,D).

**Figure 7.**
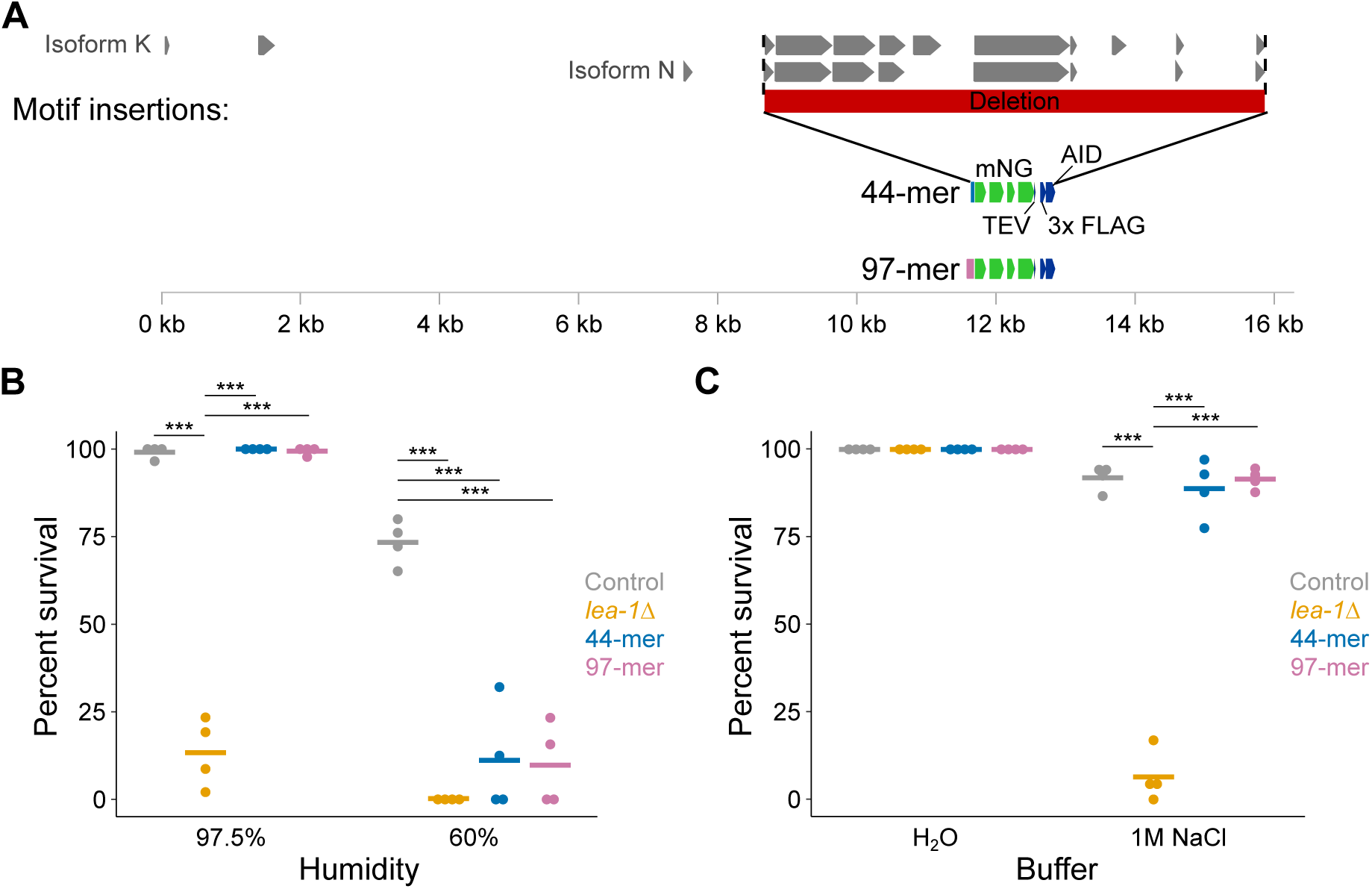
Minimal LEA-1 motifs promote desiccation and osmotic stress survival *in vivo*. **A)** Endogenous *lea-1* sequence was largely deleted and replaced with sequence encoding codon optimized, mNG-tagged, idealized motifs of 44 amino acids (4x 11-mer) or 97 amino acids. **B)** Desiccation survival of control (*daf-2*), *lea-1Δ*, 44-mer::mNG, and 97-mer::mNG animals is shown. *lea-1Δ* mutants had significantly lower survival than controls at both 97.5% RH (p=2.3×10^-6^, unpaired T-test, n=4) and 60% RH (p=4.2×10^-7^, unpaired T-test, n=4). At 97.5% RH motif-expressing worms had survival that was statistically indistinguishable from control, and significantly improved relative to *lea-1Δ* animals (44-mer p=0.36, 97-mer p=0.78, unpaired T-tests, n=4). When dehydrated at 60% RH, motif-expressing worms each had significantly reduced survival (44-mer::mNG p=2.7×10^-4^, 97-mer::mNG p=7.4×10^-5^, unpaired T-tests relative to control, n=4). **C)** *lea-1Δ* mutants, but not motif-expressing animals, were sensitive to osmotic stress in 1M NaCl for 4 hr (*lea-1Δ* p=7.5×10^-7^, 44-mer::mNG p=unpaired T-test vs. control, n=4). Experiments in B and C were conducted in dauer worms in a *daf-2(e1370)* background. *** indicates p<0.001.

Expression of either the 44 amino acid motif or 97 amino acid motif was sufficient to replace full-length LEA-1’s role in promoting desiccation tolerance of animals at 97.5% relative humidity (Fig. 7B). When exposed to 60% relative humidity, the protective capacity of these motifs was limited, and survival was reduced to levels comparable to *lea-1Δ* mutants. Expression of each motif in lieu of the complete LEA-1 protein was also sufficient to promote survival of osmotic stress in 1M NaCl (Fig. 7C). Survival of motif-expressing animals was indistinguishable from controls and significantly improved relative to *lea-1Δ* mutants. These results indicate that short repetitive motif sequences likely account for much of the functionality of the full length LEA-1 protein during osmotic stress and desiccation.

## Discussion

LEA proteins are commonly found in animals with interesting extremotolerant abilities but with limited genetic tools. Therefore, a common approach has been to study LEA proteins (and fragments thereof) in heterologous systems [20, 21, 23, 47, 48, 51–54]. While such approaches have revealed many potential functions of LEA proteins, there has been limited evidence to bridge protective abilities observed *in vitro* and in heterologous systems to the endogenous, *in vivo* context in an animal [10, 29, 55]. Therefore, we were motivated to use *C. elegans* as a model to test for *in vivo* functions of LEA-1 in its native context.

Previous studies of LEA-1 in *C. elegans* have often relied on RNAi phenotypes [10, 29]. We generated a true null mutant (*lea-1Δ*) that lacks the entire gene, in order to eliminate the possibility of residual LEA-1 expression or function. Comparison of this mutant to the insertion and deletion mutant *lea-1(tm6452)* reveals similar sensitivities to a variety of stresses, including desiccation, osmotic stress, and heat. Importantly, these phenotypes were assessed in dauer larvae, in which LEA-1 is expressed more highly at both the level of mRNA and protein [10, 35]. DUR-1, the other group 3 LEA protein of *C. elegans*, is also upregulated in dauer larvae, but is apparently unable to compensate for the loss of LEA-1 [29]. LEA-1 does not significantly impact lifespan or brood size (Fig. 3), which demonstrates the importance of the protein specifically in the context of stress resistance. Similarly, accumulation of polyglutamine protein aggregates was exacerbated by deletion of LEA-1 in dauer worms, but not in normally aging worms (Fig. 2D, Fig. S2). Effects of LEA-1 on protein aggregation have previously been demonstrated *in vitro* and in heterologous contexts [26, 37, 56]. We present evidence that LEA-1 can affect proteostasis *in vivo* in a multicellular animal. It would be fascinating to further explore any differential effects of LEA-1 with other types of aggregation-prone proteins and in tissues beyond the body wall muscle. Because LEA-1 is expressed highly in body wall muscle and is required there for desiccation survival and osmotic stress resistance, it is possible that improving proteostasis in this tissue contributes to survival of these stresses.

Adding endogenous, single-copy fluorescent tags to LEA-1 without disrupting LEA-1 function revealed the sites of its expression and *in vivo* dynamics in response to stress. In dauer larvae, mNG-tagged LEA-1 is expressed most notably in the germline, body wall muscle, pharynx, excretory cell, as well as some unidentified cells in the head. Expression patterns were similar in worms tagged with mYPET, although some sites of expression were more faint. Surprisingly, blotting for the 3x FLAG epitope included in both mNG and mYPET tags revealed significantly fewer isoforms tagged by mYPET than expected based on isoform predictions if all isoforms were expressed at high levels (Fig. 1, Fig. 4B). It is possible that the C-terminal exons are not actually included in as many isoforms as expected, or that some isoforms are expressed at low levels. The difference in the number of prominent isoforms labeled by each tag is further reflected in the significant increase in expression of mNG-labeled proteins but not mYPET- labeled proteins during desiccation (Fig. 4B-H). The upregulation of mNG-tagged proteins does not seem to be specific to a single isoform, but rather, multiple isoforms are expressed more highly during desiccation (Fig. 4B). Therefore, the increased fluorescence seen *in vivo* likely reflects the summation of these isoforms of varying sizes and compositions. LEA-1 has been reported to be regulated by transcription factors including *daf-16*/FOXO and *skn-1*/Nrf, but whether such regulation is tissue-specific or unique to some isoforms remains unexplored [57, 58]. Expression of LEA-1 in several tissues, and the upregulation of multiple isoforms led to two questions: 1) In which tissue or tissues is LEA-1 required for stress resistance? 2) Are particular isoforms more effective at conferring stress resistance?

To address the first question we utilized the auxin-inducible degradation system to deplete LEA-1 in various tissues [44]. By expressing TIR1 under the control of different promoters, we had control over the tissues in which proteins containing the auxin inducible degron (AID) tag were degraded. The AID sequence was included on the mNG tag and therefore should deplete the same isoforms containing the 3x FLAG tag (Fig. 4B). It is possible that some predicted isoforms that are not tagged with the mNG construct could escape degradation (for example isoforms C, D, and E, Fig. 1), although we do not know if such isoforms are expressed or their relative abundance. Furthermore, even though we observed decreased fluorescence in most cases (Fig. S4), depletion of AID-containing proteins may be incomplete. Incomplete degradation could result from insufficient levels of TIR1 (based on the strength of the promoter driving it) in tissues with high levels of LEA-1. For example, with *myo- 2p* driven TIR1, some expression of mNG::LEA-1 remained in the pharynx – particularly in the posterior bulb (Fig S4E). Thus, we cannot rule out the possibility that LEA-1 is required in the pharynx for survival of desiccation and osmotic stress. In contrast, depletion of LEA-1 in body wall muscle resulted in significantly reduced survival of both desiccation and osmotic stress (Fig. 5F,G). The consensus site of action from these two stresses lends confidence to the result. However, given the caveats that some untagged isoforms may remain, and that there could be incomplete degradation (even if fluorescence is depleted below detectable levels), we cannot formally conclude that LEA-1 is not required in other tissues. Rather, we present evidence that LEA-1 is required in at least body wall muscle to survive desiccation and osmotic stress. Our data suggest that other tissues can survive at least with reduced levels of the LEA-1 isoforms that are induced in dauer larvae.

To answer the second question of which LEA-1 isoforms are most functional in conferring stress resistance, we initially employed a heterologous expression approach. *E. coli* provides an ideal system to facilitate expression of proteins and rapidly screen for desiccation resistance. The fact that multiple LEA-1 isoforms could improve desiccation tolerance of bacteria turned our attention to conserved motifs contained within these isoforms. Similar motifs from LEA proteins of other organisms have been demonstrated to function in stress resistance [21, 39, 47, 48, 59]. These LEA motifs acquire secondary structure during desiccation-like conditions and specifically are thought to form amphipathic alpha-helices [19, 49, 60]. The 11- mer we identified is likely to form an amphipathic helix based on the amino acid composition and locations as depicted in Fig. 6C. Although the amino acid identity is different from other similarly identified LEA motifs, the putative structural similarities and charge distribution hint at the relative importance of structure and charge distribution over sequence identity [12, 60].

Expression of the 11-mer alone was sufficient to improve bacterial desiccation survival. A longer sequence (44-mer) marginally improved survival relative to the 11-mer (p=0.02, unpaired T-test, n=7), similar to a 97-mer (p=0.05, unpaired T-test, n=7). Thus, an increased number of motif repeats may improve survival. The importance of multiple motifs occurring in the same contiguous peptide versus the possibility of oligomerization remains unclear. Notably, in bacteria the expression levels of LEA proteins and motifs are likely much higher than in endogenous contexts.

Inserting motifs into the endogenous *lea-1* locus of *C. elegans* allowed us to test if the identified motifs are sufficient to confer desiccation *in vivo*. LEA motifs were highly effective substitutes for the full-length protein for phenotypes of osmotic stress resistance and desiccation at 97.5% RH (Fig. 7). This suggests that the function of LEA-1 can in large part be explained by the presence of these amino acid repeats. Unique combinations of motifs or longer contiguous sequences may allow for increased functionality – particularly in the context of more severe desiccation. This may further explain the apparent complexity and variety of isoforms of the protein (Fig. 1, Fig. 4B). If there is limited selective pressure for maintenance of a single long contiguous sequence, then many functional exon combinations may arise. This could contribute to the large number of proteins produced by this gene. An optimal number of motifs and the most functional combination of motif variants appended to each other remain to be characterized.

The ability of LEA motifs to function similarly in a single-celled prokaryote (*E. coli*) and a multicellular eukaryote (*C. elegans*) encourages the possibility of engineering broadly functional desicco-protectants. LEA proteins and motifs that can function in cells with fundamentally different subcellular organization likely harbor basic biochemical properties that promote cell and organismal survival. Relative to bacteria, multicellular animals face further challenges during desiccation including coordination of a response across multiple tissue types and protecting diverse organelles and subcellular compartments. *C. elegans* is a prime animal model for continued study of these *in vivo* mechanisms of desiccation tolerance.

## Conclusions

In summary, we created a true null mutant of LEA-1 to study its role *in vivo* and characterized several phenotypes of stress sensitivity. LEA-1 is expressed in multiple tissues, and it is required at high levels in body wall muscle to carry out its functions. We identified LEA- 1 motifs that are sufficient to improve bacterial desiccation survival. These motifs also function in place of full length LEA-1 to promote desiccation tolerance *in vivo*. Future work is needed to dissect the principles by which the peptide repeat sequences function. Still, the conserved function of LEA-based peptides in a single-celled prokaryote and a multicellular animal suggests the possibility of engineering LEA-1-derived peptides that broadly promote desiccation tolerance.

## Methods

### CRISPR editing in C. elegans

To generate a deletion of *lea-1* and to insert mNG and mYPET fluorescent tags into the endogenous gene locus, CRISPR methods were employed [33, 34]. Using the self-excising cassette (SEC) system, sgRNA sequences were added to the sgRNA-Cas9 plasmid pDD162 using the NEB Q5 site-directed mutagenesis kit. Homology arms of ∼500-700bp were added to plasmids carrying the SEC and repair templates. To insert a myo-2p::GFP::myo-2 3’UTR reporter to track the null deletion of *lea-1*, homology arms were cloned into plasmid pDD317. To insert mNG or mPYET tags, homology arms were cloned into pUA77 and pDD283 respectively.

To replace the endogenous *lea-1* sequence with LEA motifs, sequence was first codon-optimized for *C. elegans* and synthesized by Integrated DNA Technologies (IDT). These stretches were cloned into pUA77 with the same upstream homology arm for insertion of mNG to the endogenous locus and the same downstream homology arm for insetion of mYPET into the endogenous locus. The same sgRNAs for each of the initial mNG and mYPET insertions were used in combination to excise the *lea-1* locus.

Worms were injected with 50 ng/μL of plasmid containing the sgRNA and Cas9, and 10- 20 ng/μL plasmid containing the repair template, along with a co-injection mix [34]. 2-3 days after injection worms were treated with Hygromycin and selected for transgene-carrying rollers lacking red co-injection mix extra-chromosomal arrays. L1 worms were heat shocked at 32 °C for 5 hr to excise the SEC. Genomic edits were confirmed by visualization of fluorescent reporters and with PCR genotyping.

Construction of most TIR1-expressing strains is described in [44]. The *col-10p*::TIR1 line was generated by cloning to combine the *col-10* promoter with the TIR1 construct of pDD356 (NEB Hifi Assembly Master Mix). Plasmid pAP082 was used to express Cas9 and a sgRNA to target the chromosome I insertion site. Worms were injected with 50 ng/μL of pAP082 and 20 ng/μL plasmid containing the repair template.

### Genotyping

Template genomic DNA was create by lysing worms in lysis buffer with proteinase K. Worms were picked into 0.2mL tubes and briefly frozen at -80 °C, then heated at 65 °C for 1hr followed by 95 °C for 15 minutes. This lysate was used for PCR genotyping. Primers were designed to confirm CRISPR genome modifications and to genotype the pre-existing allele *lea- 1(tm6452)* during backcrossing. Sequences of these primers can be found in Supplemental Table 1. PCR genotyping was carried out using either Gotaq or Q5 High Fidelity Polymerase (NEB). Annealing temperature and extension time were adjusted based on the melting temperature of the primers and size of the amplicon.

### C. elegans maintenance

Worms were maintained according to standard laboratory conditions on nematode growth media (NGM) plates, fed OP50 *E. coli*, and stored at 20 °C. Standard methods were employed for crossing worms, and males were generated by keeping L4 worms at 32 °C for 5-6 hr and 25 °C overnight. The following strains and alleles were used: N2, *lea-1(tm6452)* (backcrossed 5x into N2), AM140 *rmIs132[unc-54p::Q35::YFP]*, AM141 *rmIs133[unc- 54p::Q40::YFP]*, CB1370 *daf-2(e1370)*, LP847 *lea-1Δ(cp423[myo-2p::GFP::myo-2 3’UTR])*, LP852 *daf-2(e1370);lea-1Δ(cp423[myo-2p::GFP::myo-2 3’UTR])*, LP858 *lea-1(cp431[mNG*::*3x FLAG::AID::lea-1])*, LP859 *lea-1(cp430[lea-1::mYPET::3x FLAG])*, LP860 *daf-2(e1370)*;*lea- 1(cp431[mNG*::*3x FLAG::AID::lea-1])*, LP861 *daf-2(e1370)*;*lea-1(cp430[lea-1::mYPET::3x FLAG])*, LP862 *rmIs132[unc-54p::Q35::YFP];lea-1Δ(cp423[myo-2p::GFP::myo-2 3’UTR])*, LP863 *rmIs133[unc-54p::Q40::YFP];lea-1Δ(cp423[myo-2p::GFP::myo-2 3’UTR])*, LP865 cpSi171*[vha-8p::TIR1::F2A::mTagBFP::AID*::NLS];daf-2(e1370);lea-1(cp431[mNG*::*3x FLAG::AID*::lea-1])*, LP866 cpSi172*[myo-2p::TIR1::F2A::mTagBFP::AID::NLS];daf- 2(e1370);lea-1(cp431[mNG*::*3x FLAG::AID*::lea-1])*, LP867 cpSi173*[col-10p::TIR1::F2A::mTagBFP::AID*::NLS];daf-2(e1370);lea-1(cp431[mNG*::*3x FLAG::AID::lea-1])*, LP868 cpSi174*[myo-3p::TIR1::F2A::mTagBFP::AID*::NLS];daf-2(e1370);lea-1(cp431[mNG*::*3x FLAG::AID::lea-1])*, LP875 *daf-2(e1370);lea-1(tm6452, 5x backcrossed)*, LP876 *reSi5[ges- 1p::TIR1::F2A::BFP::AID*::NLS::tbb-2 3’UTR];daf-2(e1370);lea-1(cp431[mNG*::*3x FLAG::AID::lea-1])*, LP877 *wrdSi18[mex-5p::TIR1::F2A::BFP::AID*::NLS::tbb-2 3’UTR];daf-2(e1370);lea-1(cp431[mNG*::*3x FLAG::AID::lea-1])*, LP882 *rmIs132[unc-54p::Q35::YFP]*;*daf-2(e1370);lea-1Δ(cp423[myo-2p::GFP::myo-2 3’UTR])*, LP883 *rmIs133[unc-54p::Q40::YFP];daf-2(e1370);lea-1Δ(cp423[myo-2p::GFP::myo-2 3’UTR])*, LP885 *daf-2(e1370);lea-1(cp432[lea- 1p::44-mer::mNG::3x FLAG::AID]*, LP887 *daf-2(e1370);lea-1(cp433[lea-1p::97-mer::mNG::3x FLAG::AID]*.

### C. elegans desiccation, heat, and osmotic stress

*C. elegans* were desiccated according to a previously established protocol [29, 35]. Briefly, desiccation chambers were made with varying ratios of glycerol:water in order to produce relative humidity (RH) of either 97.5% or 60%[61]. Dauer worms were formed by moving *daf-2(e1370)* mutant embryos to 25° C. Dauer worms were picked into ∼1.5-2*µ*L droplets of water and initially exposed to 97.5% for 4 days as preconditioning. Worms were either rehydrated in M9 or moved to 60% RH desiccation chambers for 1 day before recovery. Survival was scored as the percentage of worms moving or responsive to physical stimulus.

Heat stress was implemented by exposing dauer worms to 37 °C. Multiple plates containing dauer larvae were moved to a 37 °C incubator for defined periods of time. Worms were scored for survival based on movement or responsiveness to physical stimulus.

To measure osmotic stress tolerance, dauer worms were added to water or 1M NaCl in a 96-well or 24-well plate for 2 hr or 4 hr. Worms were then transferred to unseeded NGM plates and survival was scored by movement or responsiveness to physical stimulus.

### C. elegans lifespan and brood size

For lifespan assays 10 L1 worms per plate were picked to an NGM plate. For each genotype five plates of ten worms were included to start each experiment. Worms were transferred to new plates every other day until they were no longer reproductive. Worms that crawled onto the sides of the dishes or were otherwise missing were censored. Survival was scored every other day by movement or responsiveness to touch with a platinum wire. Three independent biological replicates were conducted.

To determine brood size, individual embryos were added to NGM plates. 15 worms were included to start in each of three experiments. Worms were transferred to new plates every other day. After transfer, progeny on plates were allowed to develop for ∼24 hr and were then counted. Total brood was calculated as the sum of all progeny produced by an individual worm. Worms that failed to hatch or crawled off the plate were censored.

### Polyglutamine protein aggregation

Strains AM140 and AM141 express polyglutamine tracts of 35 or 40 repeated glutamine residues fused to YFP [40]. These strains were crossed with *lea-1Δ* (LP847). To assess protein aggregation during aging the worms were picked to plates and allowed to develop for 4 or 8 days. Worms were periodically transferred to fresh plates to separate them from progeny. The number of aggregates per worm was scored by imaging worms and counting the number of fluorescent puncta in the body wall muscle of each worm.

To determine the number of aggregates in dauer worms the AM140 and AM141 reporters were crossed into *daf-2(e1370);lea-1Δ* (LP852) to put the reporter in a *daf-2* mutant background. Dauer formation was induced as before by growth at 25 °C. The number of aggregates per worm was counted after 5 days (including development and arrest as dauer larvae).

### Fluorescent imaging and quantification

Worms expressing mNG::LEA-1 or LEA-1::mYPET were imaged on a Nikon TiE stand with CSU-X1 spinning disk head (Yokogawa), 514 nm solid state laser, and ImagEM EMCCD camera (Hamamatsu). Metamorph software was used for image acquisition. To quantify fluorescence in control dauers, desiccated dauers, and dauers exposed to 1 M NaCl for 2 hr, worms were imaged with a 10x objective. Images were imported into FIJI for analysis. Whole worms were outlined manually. The total fluorescence intensity was measured. The outline of the worm was moved to background area of the image to obtain a background measurement. The fluorescence intensity was calculated by subtracting the background from that of the worm. For representative images of worms higher magnification objectives (20x or 60x) were used.

### Western blotting

Protein for western blotting was obtained from worms expressing mNG::3x FLAG::LEA-1 or LEA-1::mYPET::3x FLAG. Mixed stage populations were washed from 3-6 NGM plates, dauer worms from plates at 25 °C, and desiccated worms from plastic dishes at 97.5% RH for 4 days. Large quantities (>20,000) of synchronous worms were obtained for dauer formation and desiccation by standard hypochlorite treatment. Worms were collected, washed in M9, pelleted, and resuspended in ∼100 µL lysis buffer containing 50 mM HEPES (pH 7.4), 1mM EGTA, 1mM MgCl2, 100 mM KCl, 10% glycerol, 0.05% NP-40, DTT, and an EDTA-free protease inhibitor tablet (added fresh to 12 mL of buffer) [62]. Worms in lysis buffer were briefly frozen at -80 °C, then sonicated on ice with a Branson Sonifier 250 wand sonicator at 50% duty until no intact worms remained (2-5 minutes in individual bouts of no longer than 1-2 minutes). The concentration of protein in lysates was determined with the Biorad Protein Assay (Bradford) according to specifications, measuring the OD595 in a total volume of 1mL in cuvettes.

A 4-12% Bis-Tris NuPAGE minigel (Invitrogen), loaded with 10 µg of protein in sample buffer (3x concentrate contains 6% SDS, 240mM Tris pH 6.8, 30% glycerol, 0.04% w/v Bromophenol blue, and 50 µL 2-ME per mL) was run in 1x NuPAGE MOPS running buffer (Novex). 10 µL of Precision Plus Protein Kaleidoscope Standard (Biorad) was loaded in lanes adjacent to samples. Gels were run at 140V for 90 minutes. Protein was transferred to a PVDF membrane activated in methanol at 90V for 90 min (in NuPAGE transfer buffer, at 4 °C).

Membranes were rinsed in PBST, and blocked in PBST + 5% BSA for 1 hr with rocking. Membranes were soaked in a 1:1000 dilution of primary antibody (mouse anti-FLAG, Sigma Cat #F1804) overnight at 4 °C with rocking. Membranes were washed 3x in PBST, then soaked in a 1:5000 dilution of secondary antibody (goat anti-mouse IgG, Invitrogen Cat #31432) for 1 hr with rocking. Membranes were washed 3 times and visualized after addition of SuperSignal West Pico PLUS Chemiluminescent Substrate (Thermo Scientific).

After initial visualization, membranes were stripped and re-blotted for tubulin as a loading control. Membranes were incubated in mild stripping buffer (15 g glycine, 1 g SDS, 10 mL Tween20, bring volume to 1 L with H20, pH 2.2) for ten minutes with rocking. Buffer was replaced, and allowed to soak for another ten minutes. Membranes were then washed 3x in PBST, blocked for 1 hr in PBST with 5% BSA, and soaked in a 1:500 dilution of primary antibody (rat anti-tubulin, Invitrogen Cat #MA1-80189) overnight at 4 °C. After washing 3x in PBST, secondary antibody (goat anti-rat IgG, Thermo Fisher Cat #31470) was added in a 1:5000 dilution and membranes soaked for 1 hr with rocking and visualized as before. A single band was detected indicating antibody specificity for tubulin.

### Protein depletion with auxin-induced degradation

Worms expressing TIR1 under various promoters were grown on plates containing 1mM auxin (indole-3-acetic acid) [43]. Embryos or L1s were picked to NGM + auxin plates and allowed to develop into dauer larvae at 25 °C for at least 5 days before exposure to desiccation or osmotic stress. Degradation of LEA-1 tagged with mNG::3x FLAG::AID was assessed by imaging worms.

### Heterologous expression in bacteria

A bacterial codon-optimized version of *lea-1* isoform K was synthesized by Genewiz. Isoforms A, D, E, and F were subcloned from that construct. 30bp of homology was added by PCR to expression constructs, and they were cloned into a PCR-linearized pDest17 using NEB Hifi Assembly Master Mix. Similarly, to clone motifs, 30bp homology was added synthesized codon-optimized DNA fragments (from Integrated DNA Technologies), and they were cloned into PCR-linearized pDest17 with NEB HiFi Assembly Master Mix. GFP and truncated GFP controls were cloned by the same method, using plasmid pFCcGi as a template to obtain the GFP sequence [63]. The *C. elegans act-2* gene was also cloned as a control from cDNA. All plasmids were transformed into NEB 5-alpha cells and correct inserts were verified by sequencing.

For desiccation experiments, plasmids containing expression constructs were transformed into *E. coli* BL21 A1 (Invitrogen). Individual colonies were picked into 3 mL LB with 100 µg/mL ampicillin (Amp) and grown overnight (∼12-16 hrs) at 37 °C in a shaker incubator. Cultures were then diluted 1:20 and 0.2% L-arabinose was added to induce protein expression for 4 hr. Protein expression was confirmed with a Coomassie gel. Bacteria were pelleted and briefly boiled to generate a lysate. Protein concentration was quantified using the Biorad Protein Assay (Bradford) according to specifications. From these protein concentrations an appropriate volume was determined to load 20 ug of total protein per well. 5 µL of pageruler prestained ladder was loaded. A 4-12% BT NuPAGE minigel was run in 1x NuPAGE MOPS running buffer. The gel was stained with Coomassie and de-stained overnight. GFP-expressing bacteria provide a convenient visual confirmation of protein expression in each experiment.

To obtain equivalent numbers of starting bacteria to measure desiccation survival, OD600 of 4 hr induced cultures was measured and volumes equivalent to an OD600 of 1.5 were added to 1.5 mL Eppendorf tubes. Bacteria were washed once in 0.85% NaCl. Bacteria were then resuspended in 1 mL 0.85% NaCl. A 20 µL sample was removed and used for 10-fold dilution of the culture. A 10-fold dilution series was plated on LB + Amp agar plates by spotting 10 µL. These control plates were grown at 37 °C overnight. The remaining 980 µL supernatant was aspirated and tubes were desiccated in a speedvac overnight (∼12-16hr). Bacteria were rehydrated in 980 µL of LB + Amp and given 1-2 hr to recover before plating a 10-fold dilution series, as with controls the day before. Bacteria on plates were grown overnight at 37 °C. Survival was calculated by dividing cfu of each desiccated sample by its control.

### Motif analysis

The MEME tool of the MEME suite (http://meme-suite.org/tools/meme) was used to identify conserved motifs within LEA-1 [50]. The protein sequence of Isoform K was used as a template since it contains nearly all possible exons of the protein. Position weight matrices, consensus motifs, and distribution of motifs throughout the protein were obtained from this analysis. The pepwheel program (https://www.bioinformatics.nl/cgi-bin/emboss/pepwheel) was used to generate the diagram of the putative alpha-helical structure formed by the 11-mer. Coloration of amino acids was added by hand.

### Statistical analysis

Statistical tests were carried out as described in figure legends. Unpaired t-tests were conducted in Microsoft Excel and ANOVAs were conducted in RStudio. Lifespan analysis was conducted using GraphPad Prism.

## Availability of Data and Materials

*C. elegans* strains generated in this study will be deposited to the *Caenorhabditis Genetics Center*. Plasmids and other reagents will be made available upon request.

## Declarations

The authors declare that they have no competing interests.

---

**Figure S1.**
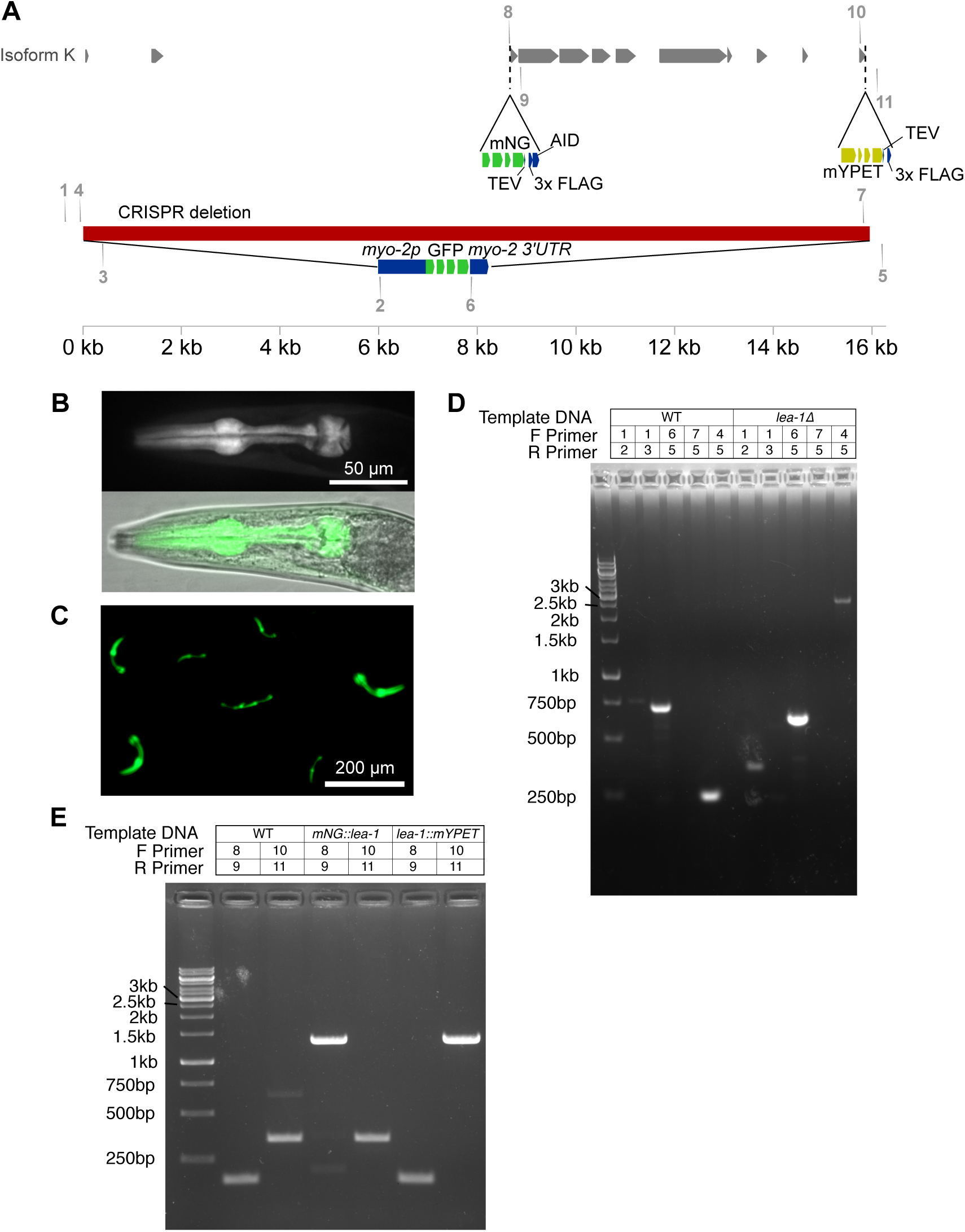
Genotyping LEA-1 alleles. **A)** A simplified schematic depicting locations of primers to genotype the various genome edits. Primers are indicated with numbers. Sequences can be found in Table S1. **B)** A representative image depicts pharyngeal GFP expression in an *lea-1Δ* mutant in which 15.8 kb of genomic sequence was replaced with *myo-2p*::GFP::*myo-2 3’UTR* as a visual marker for the deletion. **C)** A representative image shows the plate level phenotype of pharyngeal GFP expression in *lea-1Δ* mutants. **D)** Primers as indicated by numbers in Figure 1 were used to amplify genomic DNA and confirm deletion of the endogenous *lea-1* locus and insertion of *myo-2p*::GFP::*myo-2 3’UTR* in the *lea-1Δ* mutant. **E)** The indicated primers were used to amplify across the loci in genomic DNA at which the mNG and mYPET tags were inserted. Genotyping confirms insertion of the mNG::3xFLAG::AID tag in a relatively N-terminal position of *lea-1* and insertion of the mYPET::3xFLAG tag at the C-terminus of the gene.

**Figure S2.**
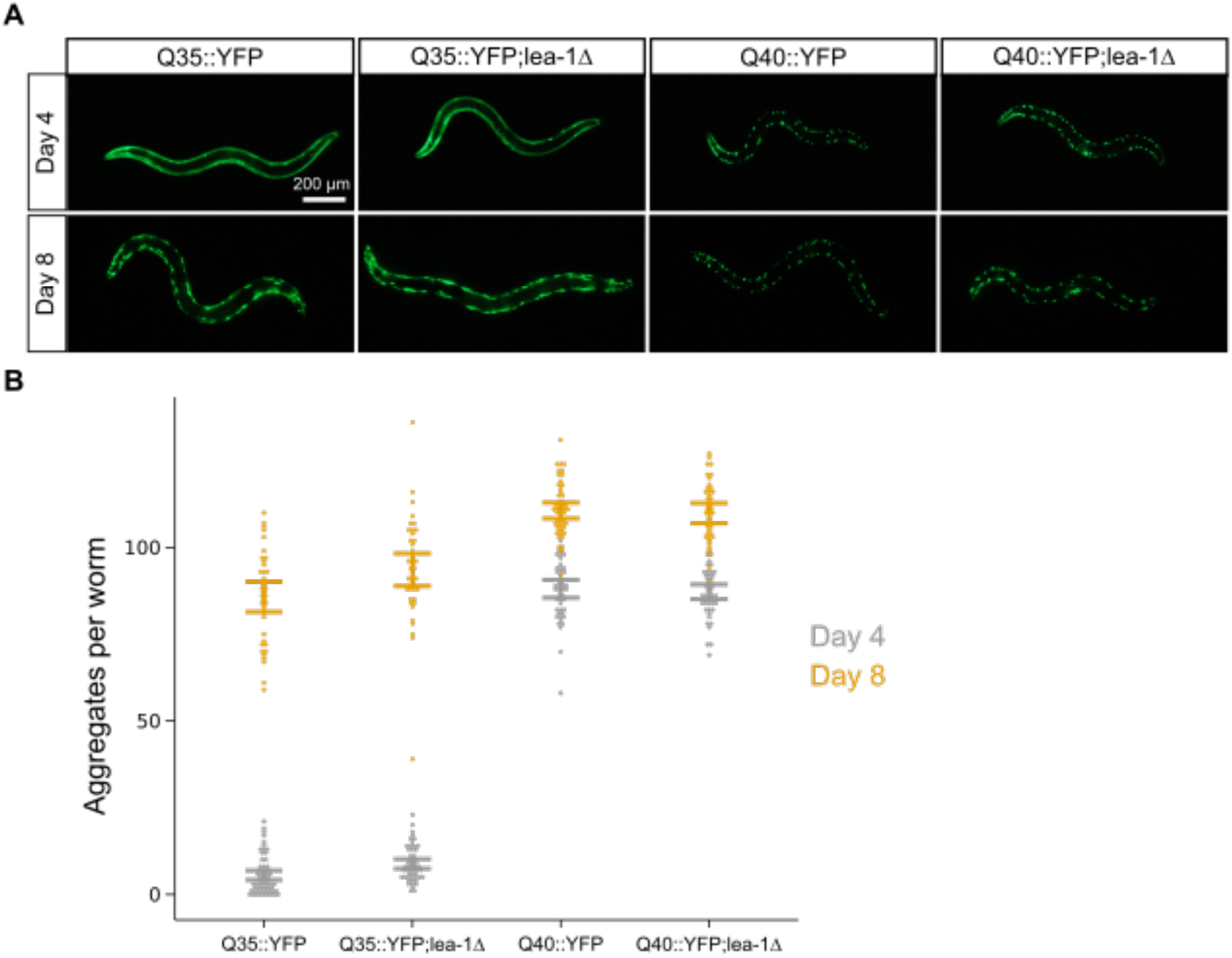
LEA-1 does not significantly alter polyglutamine protein aggregation due to age. **A)** Representative images of 4 and 8 day old worms expressing polyglutamine::YFP constructs in body wall muscle. **B)** There are no significant differences in the number of polyglutamine aggregates between control and *lea-1Δ* animals for either Q35::YFP (day 4 p=0.23, day 8 p=0.42, n=3, unpaired T-test) or Q40::YFP (day 4 p=0.81, day 8 p=0.32, n=3, unpaired T-test).

**Figure S3.**
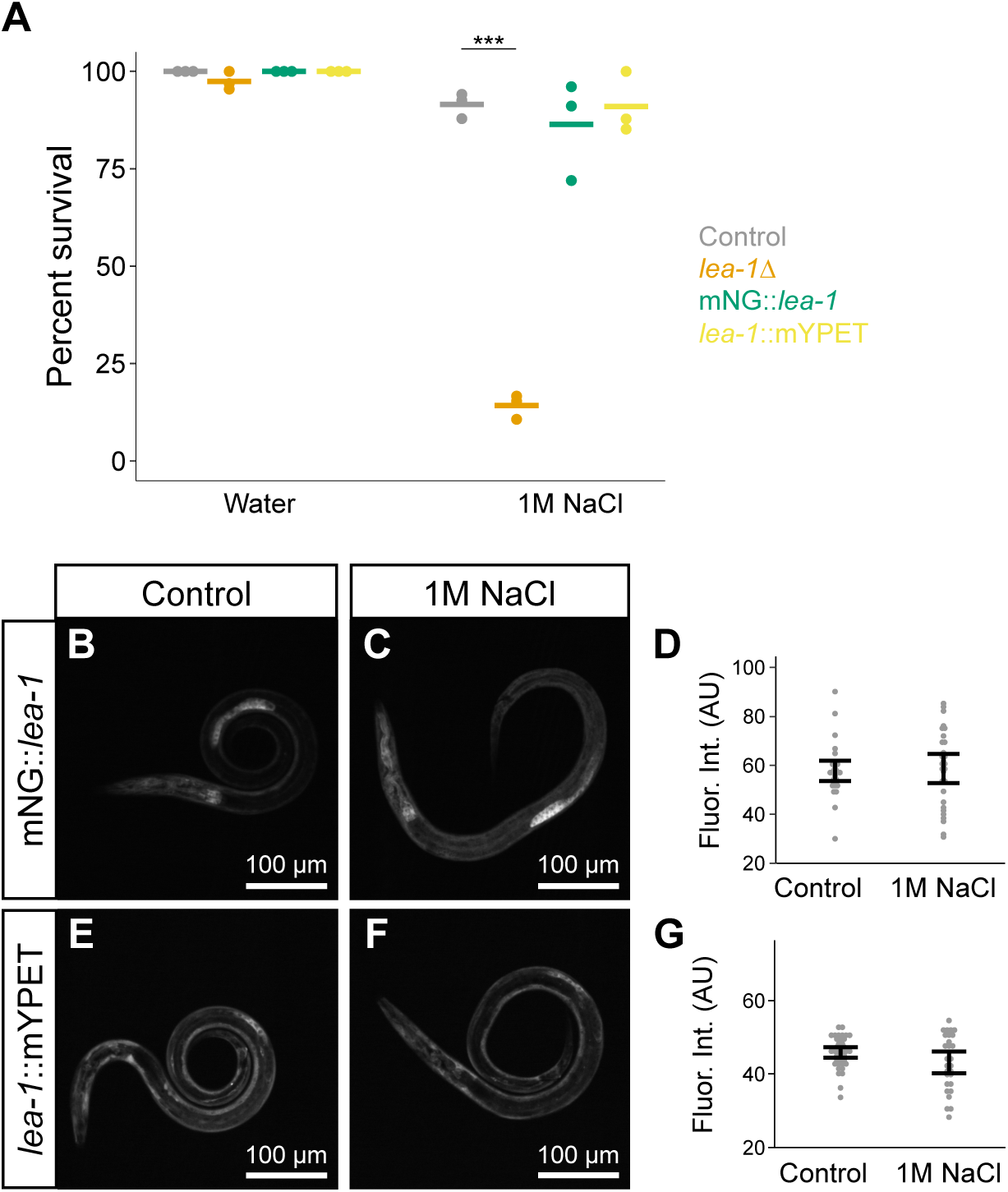
LEA-1 expression does not increase in response to short-term osmotic stress. **A)** Fluorescent tags to not disrupt function of LEA-1 during osmotic stress. Survival is plotted for worms exposed to either water or 1M NaCl for 2 hr. Bars represent mean survival. Neither mNG::*lea-1* nor *lea-1*::mYPET worms were significantly different from control worms exposed to 1M NaCl. **B)** Representative image of a control mNG::*lea-1* dauer worm. **D)** A mNG::*lea-1* dauer larvae exposed to 1M NaCl for 2 hr. **E)** mNG fluorescence is not significantly altered in worms exposed to 1M NaCl relative to controls (p=0.86, n=3 replicates, unpaired T-test). **F)** A representative control dauer worm expressing *lea-1*::mYPET. **G)** An *lea-1*::mYPET dauer after 2 hr in 1M NaCl. **H)** Fluorescent intensity is not significantly different between controls worms expressing *lea-1*::mYPET (p=0.36, n=3 replicates, unpaired T-test). All worms were in a *daf- 2(e1370)* background. *** indicates p<0.001.

**Figure S4.**
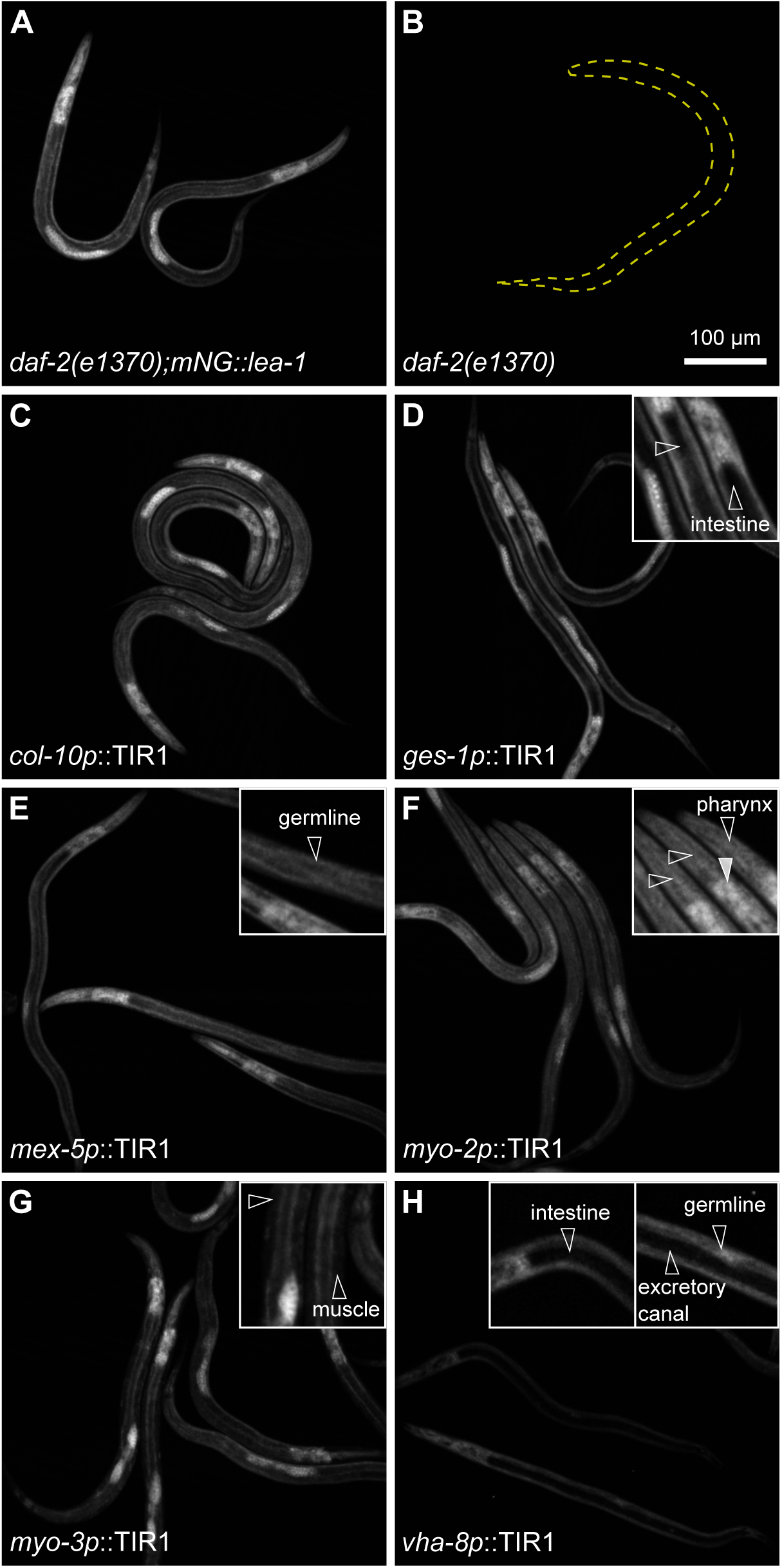
Representative images of auxin-induced depletion of mNG::*lea-1* in TIR1 expressing strains. **A)** Baseline expression of mNG::LEA-1 in the absence of TIR1 expression. **B)** *daf-2* mutant animals have no detectable baseline fluorescence. Yellow dashed lines indicate the outline of a worm. **C)** TIR1 expressed from the *col-10* promoter should deplete LEA-1 in the hypodermis. It had a minimal effect on expression levels in worms. **D)** TIR1 driven by the *ges-1* promoter reduced LEA-1 in the intestine. **E)** TIR1 under the control of the *mex-5* promoter depleted LEA-1 in the germline. Some worms retained a significantly reduced amount of germline LEA-1. **F)** *myo-2p*::TIR1 worms have reduced LEA-1 in the pharynx. This construct did not totally deplete LEA-1. In particular, expression levels are still relatively high in the posterior bulb of the pharynx (solid arrowhead). Depletion in the more anterior regions of the pharynx was more pronounced (open arrowheads). **G)** Expression of *myo-3p*::TIR1 significantly reduced LEA-1 levels in body wall muscle. **H)** TIR1 driven by the *vha-8* promoter is expressed in multiple tissues and depletes LEA-1 in the excretory cell (and canal), intestine, and germline. TIR-1 expression was also observed in hypodermis and some cells of the head. The 100 µm scale bar applies to all images. All images were taken with the same microscope settings and are shown on the same intensity scale. For the 2x magnified insets the brightness was adjusted to facilitate visualization. All worms were grown on plates containing 1mM auxin. Open arrowheads indicate sites of LEA-1 depletion in each strain.

**Figure S5.**
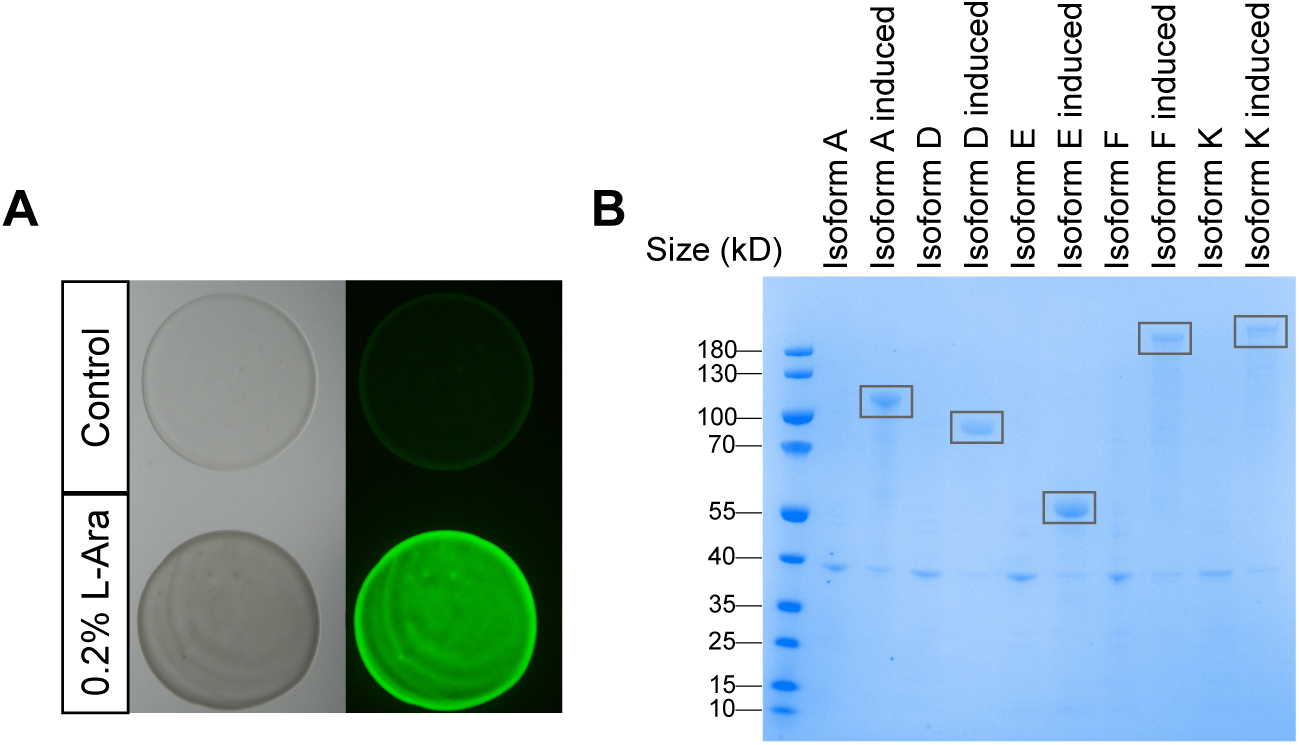
Controls for heterologous expression of proteins in bacteria. **A)** BL21 *E.coli* carrying pDest17 driving expression of GFP are induced by growth in media with 0.2% L-arabinose. Samples of bacteria from liquid cultured were spotted onto LB agar plates for imaging. **B)** A Coomassie stained 4-12% Bis-Tris gel shows expression of *C. elegans* LEA-1 isoforms A, D, E, F, and K in BL21 *E. coli* when induced with 0.2% L-arabinose. 20 µg of total protein was loaded per lane.

**Figure S6.**
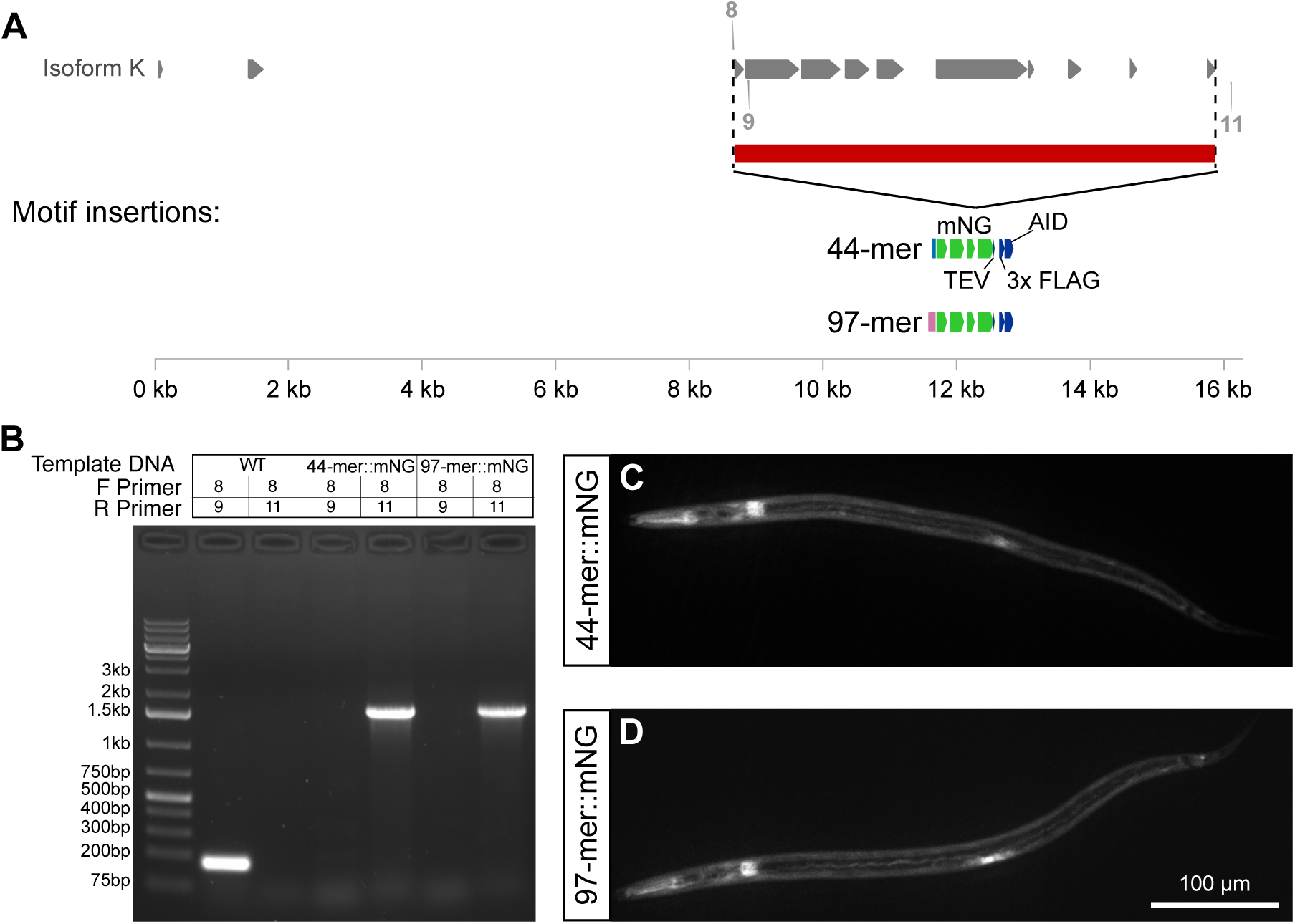
Characterization of LEA-1 motif-expressing worms. **A)** A schematic depicts the regions of genomic deletion and insertions and the primers used for genotyping these edits. **B)** PCR genotyping confirms deletion of the majority of LEA-1 exons and insertion of sequence encoding 44-mer::mNG or 97-mer::mNG. **C)** A representative image depicts *in vivo* expression of the 44-mer::mNG fusion protein in a dauer worm. **D)** A representative image depicts *in vivo* expression of the 97-mer::mNG fusion protein in a dauer worm. Worms in C and D were in a *daf-2(e1370)* background.

## Acknowledgements

We thank members of the Goldstein lab for helpful discussions and feedback. Ari Pani offered guidance on plasmid design and CRISPR editing. Priya Hibshman provided advice and assistance with Western blotting. Rob Dowen provided a plasmid from which the promoter of *col-10* was cloned. JDH was supported by the National Institutes of Health (NIH) 1F32GM131577. This work was also supported by NSF grant IOS 2028860 awarded to BG. Some strains were provided by the CGC, which is funded by NIH Office of Research Infrastructure Programs (P40 OD010440). The *tm6452* allele of *lea-1* was obtained from the National Bioresource Project in Tokyo, Japan [64].

---

**Table S1:**
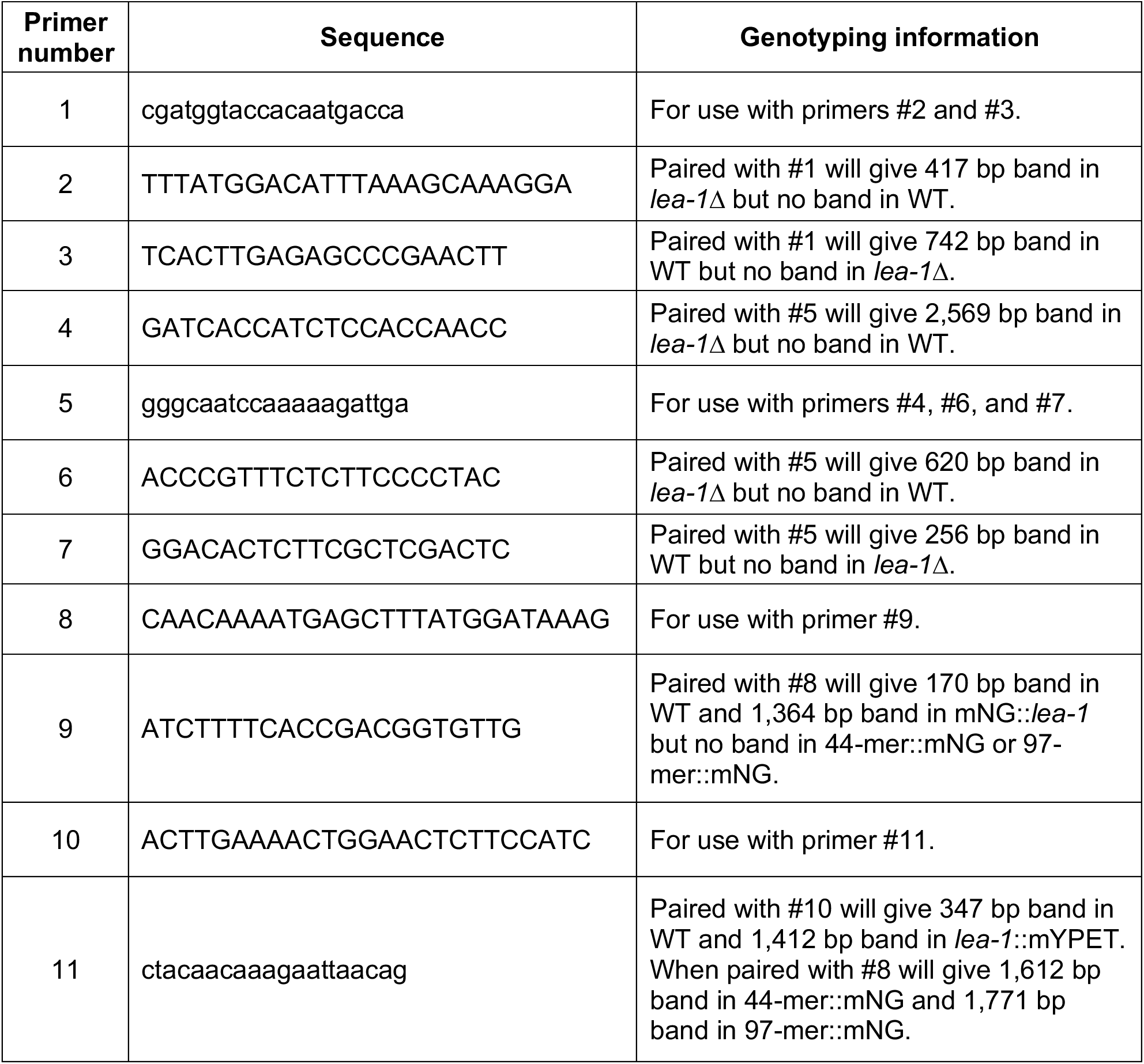
Genotyping primers for *lea-1* genomic edits.

## Notes

### Competing Interest Statement

The authors have declared no competing interest.

